# Cholesterol constrains pore formation by mammalian gasdermins

**DOI:** 10.64898/2026.10.08.757701

**Authors:** Yisha Liang, Isabella R. Evavold, Sepideh Parvanian, Makiko Hayashi, Emerson Glassey, Gema González-Rubio, Inga M. Spannaus, Victoria De Andrade, Benjamin L. Allsup, Fan Fei, Bryan D. Bryson, Aaron G. Schmidt, Tyler E. Jacks, Christopher S. Garris, Charles L. Evavold

## Abstract

Pyroptosis is a lytic form of cell death executed by gasdermin pores. Through a CRISPR screen to uncover macrophage genes required for gasdermin D (GSDMD) pore activity, we identify the endosome adaptor Rabgef1 as a positive regulator of pore formation. Rabgef1 deficiency preserves NT-GSDMD membrane binding but prevents NT-GSDMD oligomerization and lytic death. Mechanistic analyses revealed that Rabgef1 deficiency is associated with an overabundance of cholesterol in the plasma membrane. Cholesterol depletion or redistribution through multiple approaches restores GSDMD oligomerization and pyroptosis. The Rabgef1-mediated control of membrane cholesterol and pore formation was found to be a general regulator of the gasdermin family. In cancer cells, organoids, and tumors, cholesterol reduction sensitizes resistant cells to GSDME pyroptosis driven by synthetic mRNAs, chemogenetic induction, and chemotherapy treatment. These results establish membrane cholesterol content as an endogenous checkpoint for gasdermin-mediated pyroptotic cell death during the execution phase, with implications for metabolic disease and cancer immunotherapy.

## Introduction

Cell death is a defining feature of host defense and tissue homeostasis^1^. In infection and cancer, the manner in which cells die provides essential context for immune recognition^2^. Viable cells contain intracellular components that act as inflammatory mediators and antigens, and their release during lytic death initiates inflammation and primes adaptive immunity^3,4^. Thus, the molecular context of cell death determines whether the immune system interprets danger or maintains tolerance.

Among regulated death pathways, pyroptosis is an inflammatory program resulting in cellular lysis by the gasdermin family of pore-forming proteins^5^. This process is often initiated by inflammasomes, which represent the principal subcellular site of caspase-1 activation^6^. Caspase-1 cleaves gasdermin D (GSDMD) to generate an N terminal fragment (NT-GSDMD) that perforates cellular membranes^7–13^. Gasdermin pores disrupt ionic balance and enable the release of bioactive IL-1 family cytokines to coordinate local and systemic inflammation and orchestrate adaptive immune responses^9,14–16^. Other gasdermin family members, including human GSDMA, GSDMB, GSDMC, and GSDME and their mouse paralogs, execute pyroptotic death programs downstream of distinct activating signals and proteases^17^. One such example is GSDME, which causes pyroptosis downstream of caspase-3/-7 activation^18–20^. Studies to define the importance of pyroptosis in host defense have highlighted key roles of pyroptotic death processes in the control of antitumor immunity^18,20–22^, with additional maladapted roles in tissue damage, autoinflammation, and sepsis^8,17,23^. Defining means to modulate pyroptosis pathways has therefore emerged as an attractive approach to activate immune responses or minimize the impact of diverse diseases.

Current knowledge of the mechanisms of gasdermin pore formation have revealed a model whereby the cleaved protein inserts into membranes enriched in acidic phospholipids^10,13,24^. As such, several lipids have been identified as positive regulators of membrane binding and gasdermin pore formation, with notable examples being phosphatidylserine (PS), phosphoinositides (PIPs), and cardiolipin (CL)^10,13,24,25^. Despite extensive study of these positive lipid interactions, negative lipid regulation remains less understood. A prior study using atomic force microscopy showed that adding cholesterol to supported lipid bilayers composed of *E. coli* polar lipid extract reduced pore and slit formation by recombinant, cleaved human GSDMD, establishing the biophysical plausibility of sterol antagonism in a model membrane^26^. Whether cholesterol acts as a general, tunable determinant across gasdermin family members within living mammalian membranes remains unknown. Moreover, whether these threshold effects might operate across cell types and cell states to alter cell fate decisions in death signaling is wholly unstudied.

Cholesterol, the predominant sterol in mammalian membranes, maintains membrane structure and organizes signaling-competent membrane microdomains^27,28^. During infection and malignant transformation, cholesterol metabolism is frequently dysregulated^29,30^. The host modulates cholesterol localization to prevent entry or intoxication by viral and bacterial pathogens downstream of Toll-like receptor (TLR) signaling, type I and II interferon (IFN) signaling, and oxysterol production^31–33^. Moreover, cancer cells within tumors activate SREBP2-driven biosynthesis and LDLR-mediated uptake to enrich cholesterol within the tumor microenvironment that serves cancer cell-intrinsic roles in increased fitness for proliferation^34^ and decreased immune recognition^30^. Elevated cholesterol promotes myeloid cell recruitment, favors tolerogenic dendritic cell activation, and induces CD8+ T cell exhaustion^35–37^. These effects weaken antitumor immunity and favor tumor progression. Modulating cholesterol metabolism therefore provides a strategy to restore immune surveillance and sensitize cancer cells to inflammatory death.

In this study, we identify the endosome-associated adaptor Rabgef1 as a regulator of gasdermin pore formation. Loss of Rabgef1 disrupts NT-GSDMD pore assembly and protects macrophages from lytic death by increasing cholesterol at the plasma membrane. Cholesterol depletion or inhibition of cholesterol synthesis restores pore formation and pyroptosis, suggesting that sterol content of a membrane directly influences gasdermin activity. Excess cholesterol broadly suppresses pore formation by multiple gasdermin family members, whereas reducing cholesterol sensitizes cancer cells to GSDME-mediated pyroptosis. These findings reveal that cholesterol operates as an inhibitory determinant of gasdermin pore formation for the gasdermin family as a whole, and the sterol content of a given host membrane sets a threshold for how readily this pore-forming family destroys topological barriers within a living cell. The results define a metabolic checkpoint that links sterol homeostasis to inflammatory cell death and suggest combined metabolic sensitization and cell death-based immunotherapy as a rational strategy for cancer treatment.

## Results

### Endosome adaptors promote GSDMD-mediated pyroptosis

We previously conducted a genome-wide CRISPR-Cas9 screen to identify putative regulators of GSDMD pore formation^38^. From this screen, the metabolic Ragulator-Rag protein complex, through mTORC1 regulation, was identified as required for GSDMD pore formation and cell death^38,39^. To explore the potential involvement of other genes that emerged from this screen, we analyzed our hitlist with an attention to hits enriched for subcellular localization based on CellMap ID proteomic datasets^40^. This analysis uncovered an enrichment of endosome related single guide (sg)RNA in cells that were resistant to pyroptosis (Figure 1A and S1A). Pyroptosis resistance was defined as cells that were unstained by the membrane impermeable nucleic acid dye propidium iodide (PI), despite expressing high levels of NT-GSDMD fused to the blue fluorescent protein (BFP) (PI-, BFP+). The most significantly enriched sgRNAs targeting endosome adaptors in both log fold change (LFC) and p-value were to those targeting the gene Rabgef1 (Figure 1B). sgRNAs targeting other endosomal genes were identified, including the homotypic fusion and vacuole protein sorting (HOPS) and class C core endosomal vacuole tethering (CORVET) (Figure 1B). To validate our top endosome adaptor screen hits, we targeted Rabgef1 for genetic ablation. Using two independent sgRNAs and single-cell clones from each sgRNA, we ablated protein production of Rabgef1, as compared to wild-type (WT) mock transfected cells (Figure 1C). After Dox induction, the percentage of BFP-positive cells measured by flow cytometry (Figure S1B) and the bulk expression of NT-GSDMD by Western blot (Figure S1C) were comparable between Rabgef1-deficient macrophages and WT macrophages. However, Rabgef1-deficient clones displayed defects in PI staining by single cell frequency during live cell imaging (Figure 1D and 1E, Video S1, S2, S3) and LDH release (Figure 1F). Consistent with a resistance to pyroptosis in Rabgef1-deficient macrophages, cellular viability measured by normalized ATP quantification was lower in WT macrophages and remained intact in Rabgef1-deficient macrophages (Figure 1G).

**Figure 1.**
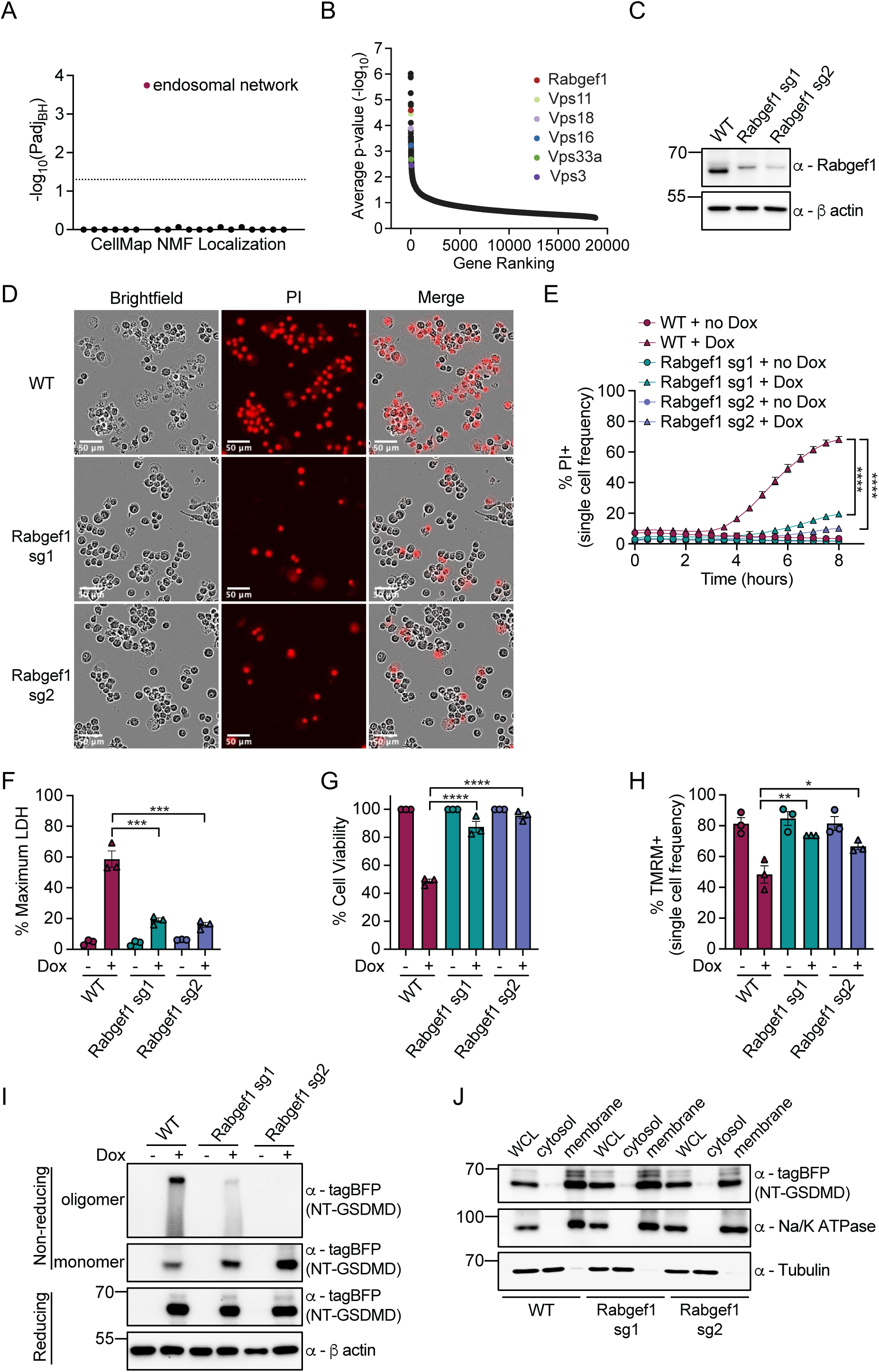
Endosome adaptor Rabgef1 promotes GSDMD-mediated pyroptosis. (A) Cell-Map NMF localization analysis of CRIPSR-Cas9 screen hits in Dox inducible NT-GSDMD expressing macrophages. (B) p-value based ranking of the hits identified in CRISPR-Cas9 screen in Dox inducible NT-GSDMD expressing macrophages. Endosome adaptors are labelled with gene symbols. (C) Western blot of Rabgef1 protein ablation in WT and Rabgef1-deficient macrophages. (D) Live cell images of WT and Rabgef1-deficient macrophages labelled with PI after Dox induction for 8 h. (E-G) WT and Rabgef1-deficient macrophages were induced with or without Dox for 8 h. Kinetics of PI positive single cell frequency to indicate membrane permeability (E), maximum LDH release to indicate membrane rupture (F) and cell viability (G) were assessed. (H) TMRM positive frequency in WT and Rabgef1-deficient macrophages with or without Dox induction for 8 h. (I) Western blot analysis of NT-GSDMD oligomers and monomers from non-reducing or reducing conditions after Dox induction for 8 h in WT and Rabgef1-deficient macrophages. (J) Western blot analysis of NT-GSDMD from whole cell lysate (WCL), cytosol and bulk membrane fractions in WT and Rabgef1-deficient macrophages after Dox induction for 8 h. Tubulin represents a cytosol marker, and Na/K+ ATPase represents a membrane marker. Data are mean ± SEM of three repeats. Immunoblots show representative result of three repeats. Statistical significance was determined by one-way ANOVA: *p < 0.05; **p < 0.01; ***p < 0.001; ****p < 0.0001.

We examined the impact of Rabgef1 deficiency on the biochemical oligomerization of NT-GSDMD. This was accomplished using an established protocol that relies on the pore conformation of NT-GSDMD being resistant to SDS solubilization when a reducing agent is not present^38^. Using this principle, we subjected WT and Rabgef1-deficient macrophages to SDS lysis after Dox induction of NT-GSDMD. When no reducing agent was added to the lysate, WT cells had mostly oligomerized NT-GSDMD (Figure 1I). In contrast, GSDMD oligomers were nearly absent from Rabgef1-deficient macrophages, yet NT-GSDMD monomeric species were abundant (Figure 1I). Treatment of the same lysates with the reducing agent β-Mercaptoethanol (BME) dissociated the oligomerized NT-GSDMD in WT cells to the monomer phase (Figure 1I).

To determine if Rabgef1 was required for membrane binding by NT-GSDMD, which is a prerequisite for oligomerization and pore formation, we conducted membrane fractionations based on centrifugation to separate the bulk membrane fraction from the cytosol fraction in WT and Rabgef1-deficient macrophages. The majority of the NT-GSDMD signal in WT and Rabgef1 deficient macrophages was in the membrane fraction (Figure 1J).

Gasdermins, including GSDMD, are known to target mitochondria to enhance cell death^19,25^. To determine the role of Rabgef1 in mitochondrial targeting by GSDMD, we fractionated mitochondria in WT and Rabgef1-deficient macrophages after NT-GSDMD expression. We found that the cytosol fraction has limited NT-GSDMD signal, as compared to the mitochondrial fraction (Figure S1E). After induction of NT-GSDMD, WT macrophages depolarized mitochondrial membranes while Rabgef1-deficient macrophages largely maintain mitochondrial potential as seen by single cell frequency of TMRM-positive cells (Figure 1H) and the mean relative TMRM intensity of all cells (Figure S1D).

These data demonstrate the endosome adaptor Rabgef1 promotes pyroptosis through oligomerization of NT-GSDMD, but Rabgef1 does not control bulk membrane and mitochondrial fraction binding of NT-GSDMD monomers.

### GEF activity, but not E3-ligase activity, of Rabgef1 promotes pyroptosis

Rabgef1 is a guanine-exchange factor (GEF) that swaps guanosine diphosphate (GDP) for guanosine triphosphate (GTP) to promote endosome trafficking via conformational change of Rab endosome adaptors, such as Rab5^41^ (Figure 2A and S2A). To determine if Rabgef1 deficiency affects the endosomal activities of macrophages, we first assessed phagocytosis using fluorescent zymosan bioparticles. We noted that phagocytosis of fluorescent zymosan bioparticles is comparable between WT and Rabgef1-deficient macrophages (Figures S2B and S2C). We next evaluated fluid phase endocytosis (pinocytosis) by measuring fluorescent Dextran uptake. While Rabgef1-deficient macrophages showed a slight defect in fluorescent intensity, overall frequency of dextran internalization was not significantly impaired (Figure S2D and S2E). We finally tested if Rabgef1 deficiency could affect LDLR-mediated endocytosis of fluorescent low-density lipoprotein (LDL). WT and Rabgef1-deficient macrophages exhibited comparable LDL internalization and fluorescence intensity (Figure S2F and S2G). Taken together, these data suggest that Rabgef1 deficiency has a limited effect on several distinct internalization pathways in macrophages.

**Figure 2.**
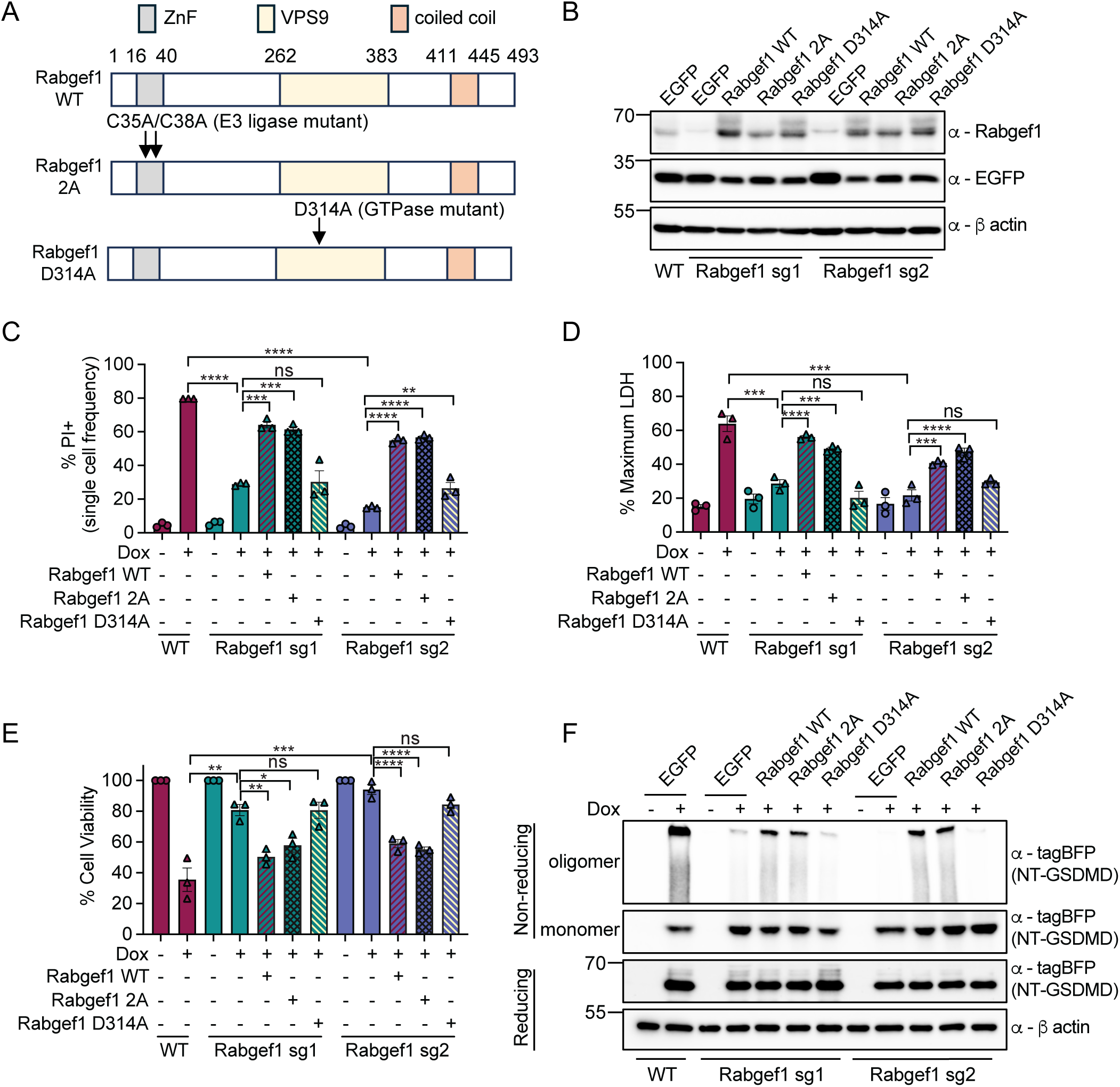
GEF activity but not E3 ligase activity of Rabgef1 promotes pyroptosis. (A) Schematic of Rabgef1 variants used in this study. Arrows indicate the point mutations of Rabgef1 with E3 ligase null mutation and GTPase null mutation. (B) Western blot analysis of protein expression of Rabgef1 variants in indicated macrophage genotypes. (C-E) The indicated macrophage genotypes were induced with or without Dox for 8 h. PI positive single cell frequency (C), bulk LDH release (D) and cell viability (E) were determined. (F) Western blot analysis of NT-GSDMD oligomers and monomers under non-reducing and reducing conditions from the indicated macrophage genotypes with or without Dox induction for 8 h. Data are mean ± SEM of three repeats. Immunoblots show representative result of three repeats. Statistical significance was determined by one-way ANOVA and unpaired two-tailed t-test: *p < 0.05; **p < 0.01; ***p < 0.001; ****p < 0.0001.

Beyond the canonical GEF function, Rabgef1 also has a described function as a non-canonical E3 ubiquitin ligase^42^ (Figure 2A and S2A). To identify functional domains within Rabgef1 that regulate GSDMD pore formation, we constructed lentiviral vectors that encode for different variants of Rabgef1. Specifically, we used the Rabgef1 WT variant, an E3-ligase null variant with two cysteine-to-alanine mutations (C35A/C38A) in the N terminal Zinc nuclease Finger (ZnF) domain hereafter referred to as Rabgef1 2A^43^, and a GEF activity null variant with a single aspartate-to-alanine mutation in the VPS9 domain hereafter referred to as Rabgef1 D314A^44^ (Figure 2A). We reconstituted Rabgef1-deficient macrophages with these different coding variants of Rabgef1 with lentivirus containing a T2A-EGFP to report on the expression of Rabgef1 and sort cells by flow cytometry with similar EGFP fluorescence. Similar expression of EGFP and Rabgef1 variants in Rabgef1-deficient macrophages was confirmed by Western blot (Figure 2B). As expected, WT cells expressing only an EGFP reporter display robust GSDMD plasma membrane pore formation and pyroptotic lysis after NT-GSDMD induction, whereas Rabgef1-deficient macrophages expressing only an EGFP reporter were defective for these activities (Figure 2C and 2D). Reconstitution of Rabgef1-deficient macrophages with Rabgef1 WT or the E3-ligase activity null Rabgef1 2A variant rescued GSDMD pore formation (Figure 2C) and pyroptotic lysis (Figure 2D) after NT-GSDMD induction. In contrast, reconstitution of Rabgef1-deficient cells with the GEF activity null Rabgef1 D314A variant did not rescue GSDMD pore formation (Figure 2C) and pyroptotic lysis (Figure 2D). Macrophages expressing WT and Rabgef1 2A variants demonstrated a decrease in cellular ATP similar to EGFP expressing WT macrophages, whereas cells reconstituted with the Rabgef1 D314A variant maintained cellular viability similar to EGFP expressing Rabgef1-deficient macrophages (Figure 2E). Furthermore, biochemical oligomerization of NT-GSDMD was rescued by expression of WT and 2A variants of Rabgef1, but not by expression of the D314A variant of Rabgef1 (Figure 2F). These data together suggest that Rabgef1 GEF activity, but not E3-ligase activity, is required for its pyroptosis-promoting activities within macrophages.

### Rabgef1 controls cholesterol content in macrophages

In humans, rare patient populations have been identified with loss-of-function alleles in genes encoding Vps family proteins of the HOPS/CORVET endosome sorting complexes^45^. These patients demonstrate developmental defects and short life expectancy^45^. Primary human fibroblasts from these patients demonstrate hallmarks of lysosomal storage defects (LSDs) including elevated staining for cholesterol content^45^. These clinical data suggested a connection between the endosome adaptor hits from our genome-wide screen and cholesterol. To assess this link directly, we extracted non-esterified free cholesterol from macrophages and quantified it with an Amplex Red assay (Figure 3A). Consistent with prior work in genome edited Hela cells^46^ and patient fibroblasts^45^, we found that Rabgef1-deficient murine macrophages contained elevated cholesterol compared to WT macrophages. Additionally, staining of total cellular cholesterol within Rabgef1-deficient macrophages with the filipin reagent demonstrates elevated mean fluorescence intensity (MFI) of single cells compared to WT cells by flow cytometry (Figure 3B).

**Figure 3.**
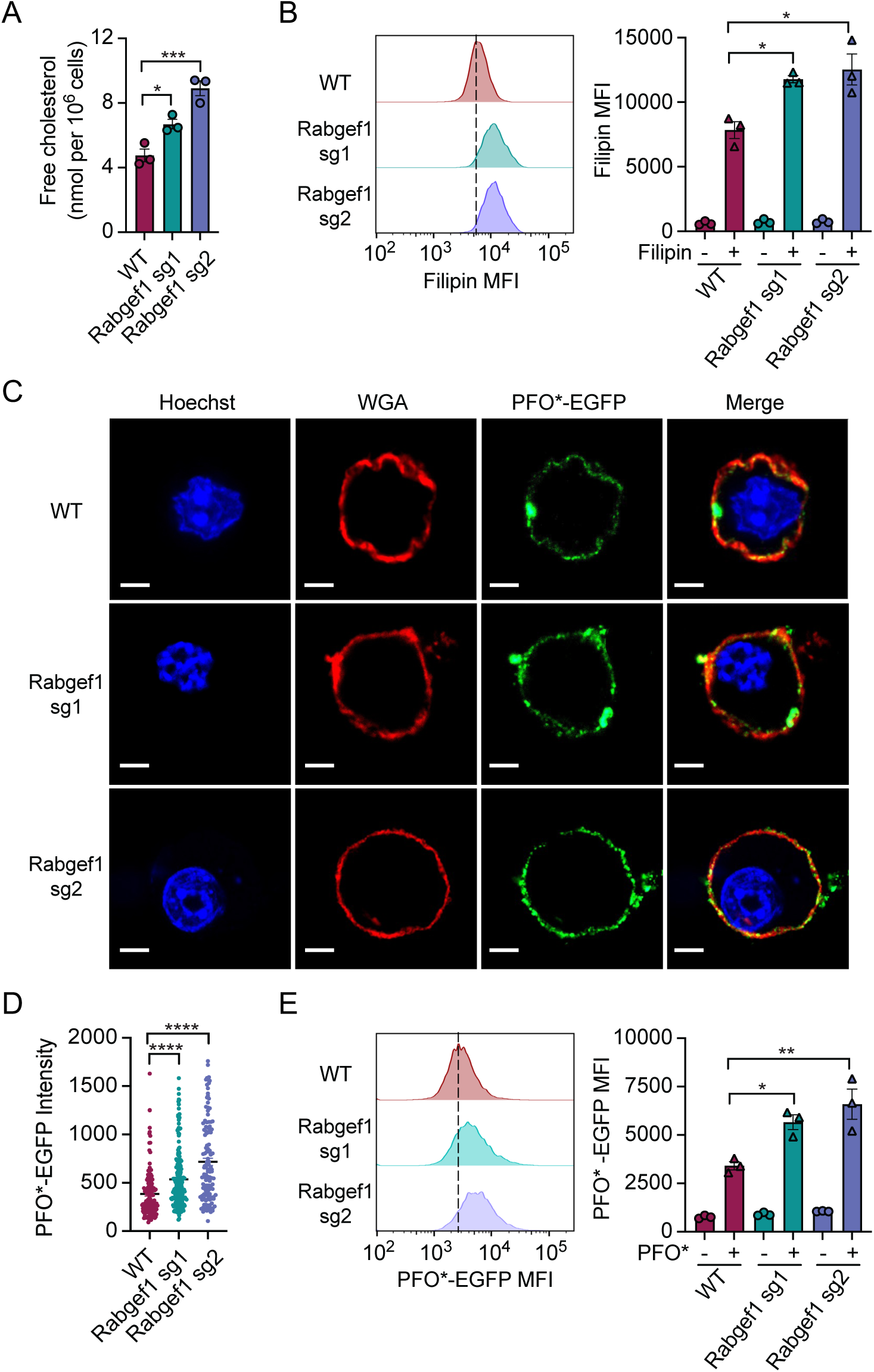
Rabgef1 deficiency results in elevated cholesterol content in macrophages. (A) Free cholesterol quantification in WT and Rabgef1-deficient macrophages. (B) WT and Rabgef1-deficient macrophages were stained with or without Filipin. Flow cytometry histograms and mean fluorescent intensity (MFI) of Filipin were analyzed. (C) Confocal images in WT and Rabgef1-deficient macrophages stained with Hoechst (nuclear, blue), WGA (cell plasma membrane, red) and PFO*-EGFP (cell plasm membrane cholesterol, green). Scale bar indicates 2 µm. (D) Violin plots of PFO*-EGFP fluorescent intensity quantified from C (n=143 for WT macrophages, n=198 for Rabgef1 sg1 macrophages, n=127 for Rabgef1 sg2 macrophages). (E) WT and Rabgef1-deficient macrophages were stained with PFO*-EGFP. Data in A, B and E are mean ± SEM of three repeats. Images in B are representative of two repeats and data in D is mean ± SEM of two independent repeats. Statistical significance was determined by one-way ANOVA: *p < 0.05; **p < 0.01; ***p < 0.001; ****p < 0.0001.

Cholesterol synthesis is transcriptionally regulated by the activity of the transcription factor SREBP2^47^. SREBP2 is an endoplasmic reticulum (ER) membrane integrated protein that is cleaved and released to the nucleus when the cell senses low ER membrane cholesterol concentrations^48^. SREBP2 activates the transcription of genes involved in uptake of cholesterol from exogenous sources, such as LDLR, and endogenous biosynthesis, such as the upstream enzyme 3-hydroxy-3-methylglutaryl coenzyme A reductase (HMGCR) as well as rate limiting enzymes including squalene epoxidase (SQLE) and 7-dehydrocholesterol reductase (DHCR7)^49^. LDLR-mediated endocytosis remained unaffected by Rabgef1 deficiency (Figure S2F and S2G), which prompted an in-depth analysis of endogenous cholesterol biosynthesis. We investigated the *de novo* cholesterol biosynthesis pathway by quantitative reverse-transcriptase PCR (qPCR) using primer probes for LDLR, SREBP2, SQLE, and DHCR7. As expected from our functional datasets on LDL internalization, the mRNA levels of LDLR were similar between WT and Rabgef1-deficient macrophages in either lipid containing (complete) or lipid-depleted (delipidated) media conditions (Figure S3A). A slight but insignificant upregulation of LDLR mRNA was noted between complete and delipidated culture conditions within a genotype (Figure S3A). Similar to a prior study that uncovered endosome adaptors as hits for a SREBP2 transcriptional reporter screen in starved conditions^46^, we observed elevation of mRNAs encoding for the transcription factors SREBP2, SQLE and DHCR7 in WT iBMDMs cultured in delipidated conditions compared to complete media (Figures S3B-D). This same elevation is observed when comparing WT versus Rabgef1-deficient macrophages in complete media, suggesting that Rabgef1-deficient macrophages turn on a SREBP2-dependent program even in fed conditions (Figures S3B-D). We observed no further enhancement to SREBP2, SQLE, and DHCR7 in Rabgef1-deficient macrophages in delipidated media (Figures S3B-D).

These findings suggest that Rabgef1 deficiency triggers an SREBP2-driven cholesterol biosynthetic program, even in cells that are normally sterol-replete via exogenous uptake. Based on these results, together with our finding that LDLR-uptake remains intact, our model supports the interpretation that Rabgef1 deficient cells accumulate cholesterol from both de novo synthesis plus uptake from outside the cell. We propose that defective endosomal trafficking disrupts the feedback that normally suppresses SREBP2 activity in cholesterol-replete cells, allowing these two sources of cholesterol to combine and expand the plasma membrane pool.

As GSDMD pores form at the plasma membrane to mediate membrane permeability and cell death^7–12^, we endeavored to measure the plasma membrane specific pool of cholesterol within single cells. Some bacteria-derived toxins, known as cholesterol-dependent cytolysins (CDCs), bind to cholesterol to mediate insertion, oligomerization, and perforation of host plasma membranes^50^. A canonical bacterial CDC pore-forming protein is Perfringolysin O (PFO) from

#### Clostridium perfringens^51^

While PFO monomers bind to plasma membrane cholesterol, binding at 37°C results in pore oligomerization and cell death^51^. Monomer-locked variants (PFO*) have been identified, through point mutations (A71T, C459A, and Y181A), which allows for monomer staining of plasma membrane cholesterol at 4°C without membrane permeabilization^52^. We purified recombinant PFO* with a SpyTag and recombinant EGFP with a SpyCatcher from *E. coli* and covalently linked these proteins by combining SpyTag with SpyCatcher to yield a highly pure and fluorescent PFO*-EGFP reagent (Figures S3E-S3H). We stained WT and Rabgef1-deficient macrophages with wheat germ agglutinin (WGA) that binds to glycans as a plasma membrane stain and Hoechst as a nuclear stain to identify single cells by fixed cell confocal microscopy (Figure 3C). Staining WT and Rabgef1-deficient macrophages with PFO*-EGFP demonstrates plasma membrane topology-specific staining for cholesterol with colocalized wheat germ agglutinin (WGA) in internal z-plane by confocal microscopy (Figure 3C).

Quantification of PM cholesterol of single cells by PFO*-EGFP MFI through confocal microscopy (Figure 3D) and flow cytometry (Figure 3E) confirms PM cholesterol is elevated in Rabgef1-deficient macrophages compared to WT macrophages. These collective results indicate that Rabgef1 is required to maintain low levels of cholesterol in the membranes of macrophages.

### Acute plasma membrane cholesterol depletion promotes GSDMD pores and pyroptosis

To determine if Rabgef1 control of pyroptosis and cholesterol content are causally linked, we sought to manipulate plasma membrane pools of cholesterol during pyroptosis. Methyl-β-cyclodextrin (MCD) is a reagent known to extract cholesterol from membranes without the need for active transporters^53^. Acute treatment of WT and Rabgef1-deficient macrophages with MCD results in decreased free cholesterol quantification, and the elevated cholesterol noted in Rabgef1-deficient macrophages is reduced by MCD treatment to a similar quantity as untreated WT macrophages (Figure 4A). Acute MCD treatment specifically depleted the PM pool of cholesterol in WT and Rabgef1-deficient macrophages, as assessed by PFO*-EGFP staining (Figure 4B). Acute MCD treatment following 7 hours of NT-GSDMD induction rescues the membrane permeability pyroptosis defects associated with Rabgef1 deficiency (Figure 4C and 4D). MCD treatment after NT-GSDMD induction also rescued the defect of Rabgef1-deficient macrophages for NT-GSDMD induced ATP loss (Figure 4E). MCD treatment in the absence of NT-GSDMD (no Dox) does not result in increased membrane permeability or lysis (Figure 4C and 4D) or a decrease in ATP content (Figure 4E), demonstrating that acute MCD treatment alone does not cause non-specific toxicity. MCD treatment rescued GSDMD oligomerization in Rabgef1-deficient macrophages (Figure 4F).

**Figure 4.**
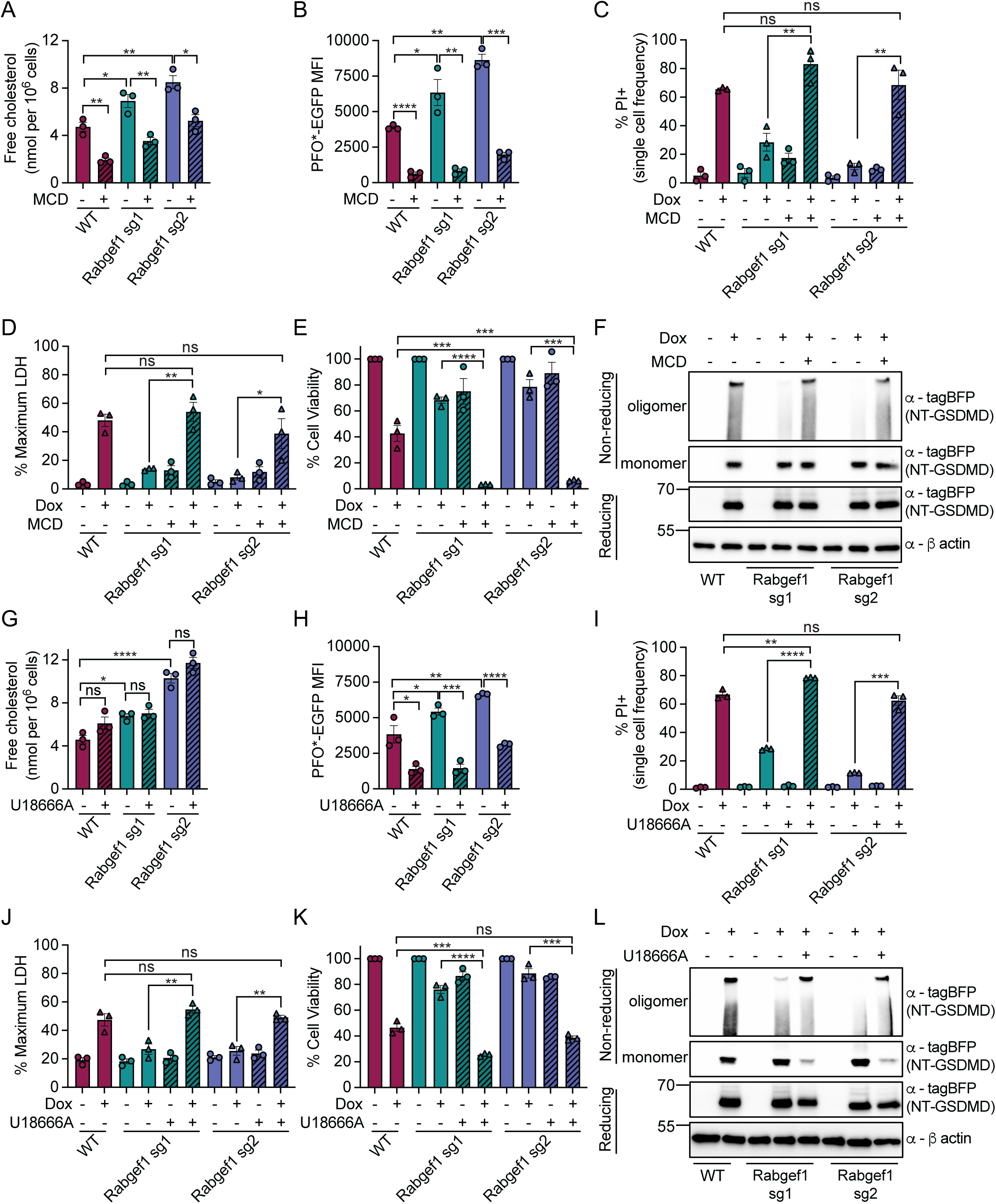
Acute plasma membrane cholesterol depletion promotes GSDMD pore formation and pyroptosis. (A) WT and Rabgef1-deficient macrophages were treated with or without MCD (10 mM) for 1 h. Free cholesterol was quantified. (B) WT and Rabgef1-deficient macrophages were treated with or without MCD (10 mM) for 1 h, followed by PFO*-EGFP staining. MFI of PFO*-EGFP to indicate cell plasma membrane cholesterol content was analyzed. (C-E) WT and Rabgef1-deficient macrophages were induced with or without Dox for 7 h, followed by treatment with or without MCD (10 mM) for 1 h. PI positive single cell frequency (C), bulk LDH release (D) and cell viability (E) were determined. (F) Western blot analysis of NT-GSDMD oligomers and monomers under non-reducing and reducing conditions in WT and Rabgef1-deficient macrophages induced with or without Dox for 7 h, followed by treatment with or without MCD (10 mM) for 1 h. (G) WT and Rabgef1-deficient macrophages were treated with or without U18666A (2 µM) for 9 h. Free cholesterol was quantified. (H) WT and Rabgef1-deficient macrophages were treated with or without U18666A (2 µM) for 9 h, followed by PFO*-EGFP staining. MFI of PFO*-EGFP to indicate cell plasma membrane cholesterol content was analyzed. (I-K) WT and Rabgef1-deficient macrophages were pre-treated with or without U18666A (2 µM) for 1h, followed by induction with or without Dox for 8 h. PI positive single cell frequency (I), bulk LDH release (J) and cell viability (K) were determined. (L) Western blot analysis of NT-GSDMD oligomers and monomers under non-reducing and reducing conditions in WT and Rabgef1-deficient macrophages pre-treated with or without U18666A (2 µM) for 1 h, followed by induction with or without Dox for 8 h. Data are mean ± SEM of three repeats. Immunoblots show representative result of three repeats. Statistical significance was determined by one-way ANOVA and unpaired two-tailed t-test: *p < 0.05; **p < 0.01; ***p < 0.001; ****p < 0.0001.

Transport of endocytosed cholesterol from lysosomes to plasma membrane relies on the Niemann-Pick disease type C1 (NPC1) protein^54^. Loss of NPC1 blocks cholesterol transport, leading to cholesterol accumulation within the lumen of lysosomes^54^. To test whether transient NPC1 inhibition would lower PM cholesterol content and rescue pyroptosis in Rabgef1-deficient cells, WT and Rabgef1-deficient macrophages were pre-treated with U18666A, an NPC1 inhibitor, followed by Dox induction of NT-GSDMD expression. We found that NPC1 inhibition has no effect on the total cellular free cholesterol concentration in either WT or Rabgef1-deficient macrophages (Figure 4G), but NPC1 inhibition significantly reduced plasma membrane cholesterol indicated by reduced PFO*-EGFP MFI (Figure 4H). NPC1 inhibition rescued membrane permeability (Figure 4I) and pyroptotic lysis (Figure 4J) in Rabgef1-deficient macrophages. Consistent with the rescued pyroptosis in Rabgef1-deficient macrophages, NPC1 inhibition also results in decreased ATP content in Rabgef1-deficient macrophages to levels comparable with WT macrophages (Figure 4K), and NPC1 inhibition rescued NT-GSDMD oligomerization in Rabgef1-deficient macrophages (Figure 4L). Taken together these data demonstrate cholesterol, specifically localized to plasma membrane, determines GSDMD pore forming activity.

### Cholesterol biosynthesis inhibitors sensitize cells to GSDMD pores and pyroptosis

We next sought pharmacologic means to manipulate cholesterol levels within our macrophages through targeting nodes in the cholesterol biosynthesis pathway (Figure 5A). WT and Rabgef1-deficient macrophages were plated in delipidated media to restrict uptake from exogenous sources to focus our attention on biosynthesis as a source of cellular cholesterol. WT and Rabgef1-deficient macrophages treated with the SREBP2 inhibitor Fatostatin display diminished PM cholesterol by PFO*-EGFP staining (Figure 5B). Rabgef1-deficient macrophages treated with Fatostatin rescue GSDMD-mediated plasma membrane permeability (Figure 5C) and pyroptotic lysis (Figure 5D). As targeting SREBP2 affects many steps within the mevalonate pathway some of which result in cellular changes unrelated to cholesterol, we sought to target late-stage rate limiting enzymes within commitment steps that are specific for cholesterol synthesis. We use the inhibitor NB598 to target the conversion of squalene to epoxysqualene by the enzyme SQLE (also known as squalene monooxygenase). WT and Rabgef1-deficient macrophages treated with NB598 display diminished PM cholesterol by PFO*-EGFP staining (Figure 5E). Rabgef1-deficient macrophages treated with NB598 rescue GSDMD-mediated plasma membrane permeabilization (Figure 5F) and pyroptotic lysis (Figure 5G). We also target the final commitment step for *bona fide* cholesterol synthesis through the conversion of 7-dehydrocholesterol to cholesterol with the small molecule AY9944 that targets DHCR7. WT and Rabgef1-deficient macrophages treated with AY9944 display diminished PM cholesterol by PFO*-EGFP staining (Figure 5H). Rabgef1-deficient macrophages treated with AY9944 rescue GSDMD-mediated plasma membrane permeability (Figure 5I) and pyroptotic lysis (Figure 5J). Consistent with functional rescue of GSDMD pores and pyroptosis, treatment of Rabgef1-deficient macrophages expressing NT-GSDMD with SREBP2 inhibitor Fatostatin, SQLE inhibitor NB598, and DHCR7 inhibitor AY9944 rescues GSDMD biochemical oligomerization (Figure 5K).

**Figure 5.**
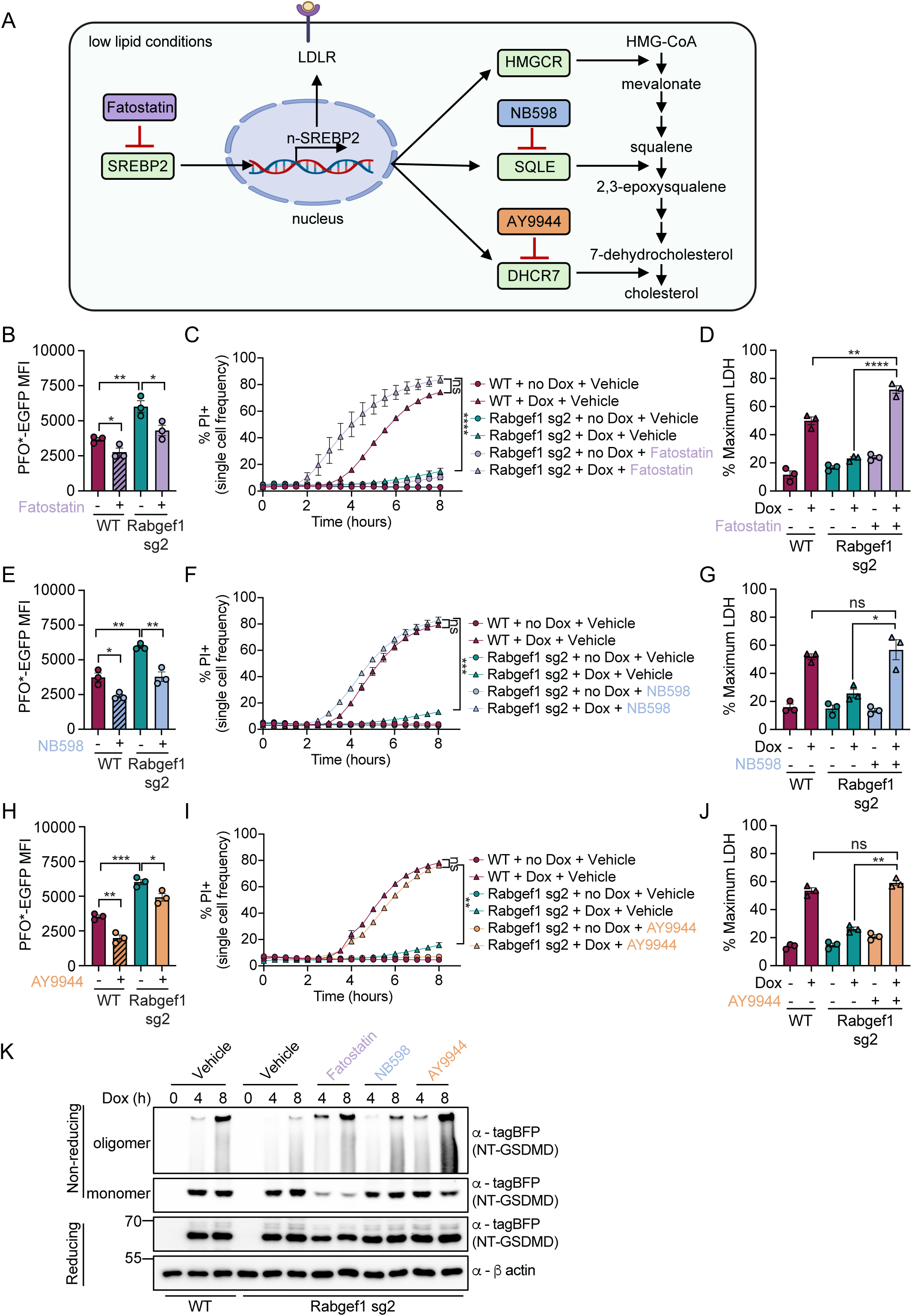
Inhibiting cholesterol biosynthesis rescues GSDMD pore formation and pyroptosis. (A) Schematic of cholesterol biosynthesis pathway and inhibitors targeting SREBP2, SQLE and DHCR7. (B, E and H) WT and Rabgef1 sg2 macrophages were treated with or without Fatostatin (20 µM) (B), NB598 (2.5 µM) (E) and AY9944 (10 µM) (H) for 9 h, followed by PFO*-EGFP staining. The MFI of PFO*-EGFP was analyzed. (C and D) WT and Rabgef1 sg2 macrophages were pre-treated with or without Fatostatin (20 µM) for 1 h, followed with or without Dox induction for 8 h. Kinetics of PI positive single cell frequency (C) and bulk LDH release (D) were determined. (F and G) WT and Rabgef1 sg2 macrophages were pre-treated with or without NB598 (2.5 µM) for 1 h, followed with or without Dox induction for 8 h. Kinetics of PI positive single cell frequency (F) and bulk LDH release (G) were determined. (I and J) WT and Rabgef1 sg2 macrophages were pre-treated with or without AY9944 (10 µM) for 1 h, followed with or without Dox induction for 8 h. Kinetics of PI positive single cell frequency (I) and bulk LDH release (J) were determined. (K) WT and Rabgef1 sg2 macrophages were pre-treated with or without Fatostatin (20 µM), NB598 (2.5 µM) and AY9944 (10 µM) for 1h, followed with or without Dox induction for indicated time points. NT-GSDMD oligomers and monomers under non-reducing and reducing conditions were analyzed by western blot. Data are mean ± SEM of three repeats. Immunoblots show representative result of three repeats. Statistical significance was determined by one-way ANOVA and unpaired two-tailed t-test: *p < 0.05; **p < 0.01; ***p < 0.001; ****p < 0.0001.

Plasma membrane cholesterol levels in epithelial and myeloid cells are dynamically regulated during immune responses to pathogens downstream of interferon (IFN) and toll-like receptor (TLR) signaling^31,32^. A major axis by which IFN and TLR signaling causes depletion of plasma membrane pools of cholesterol is through cholesterol relocalization to esterified pools in lipid droplets, downstream of oxysterol production and signaling. A prior publication noted that membrane permeability increased after treatment with the bioactive oxysterol 25-hydroxycholesterol (25HC) but not a related oxysterol 19-hydroxycholesterol (19HC) downstream of non-CDC bacterial pore forming toxin treatment and pyroptotic NLRP3-inflammasome signaling^55^. WT and Rabgef1-deficient macrophages treated with 25HC display diminished PM cholesterol by PFO*-EGFP staining that does not occur with 19HC treatment (Figure S4A). Rabgef1-deficient macrophages treated with 25HC, but not 19HC, rescue GSDMD-mediated plasma membrane permeability with faster kinetics than WT cells (Figure S4B). Furthermore, treatment of Rabgef1-deficient cells with 25HC rescues pyroptotic lysis (Figure S4C) and reduces cell viability (Figure S4D). Consistent with functional rescue of GSDMD pores and pyroptosis, treatment of Rabgef1-deficient macrophages expressing NT-GSDMD with 25HC rescues GSDMD biochemical oligomerization (Figure S4E).

### Mammalian gasdermin pore-forming domains are controlled by cholesterol content

We sought to determine whether Rabgef1 and cholesterol play a general role in regulating pyroptosis induced by other gasdermin family members. Toward this end, we utilized modified base containing mRNAs, which enable genetic payload expression in experimental and clinical contexts. Commonly employed base modifications within mRNAs include pseudo-uridine as in the FDA-approved COVID vaccines^56,57^ as well as 5-methoxy-uridine (5-mo-U) and 5-methyl-cytosine (5-m-C), which results in diminished immune recognition by the pattern recognition receptors MDA5 and RIG-I, endosomal Toll-like receptors (TLRs), and protein kinase R (PKR)^58^. We generated a panel of modular gasdermin variants as synthetic mRNAs with 5-mo-U and 5-m-C substitutions. This “synthetic killer mRNA” panel consists of the full-length (FL) inactive forms, and the N terminal (NT) pore forming domains of human gasdermin A (GSDMA), mouse gasdermin A3 (GSDMA3), human gasdermin B (GSDMB), human gasdermin C (GSDMC), human gasdermin D (GSDMD), and human gasdermin E (GSDME) (Figure S5A). We also generate EGFP modified mRNA as a reporter and carrier mRNA to monitor transfection efficiency into various cell types after co-encapsulation of gasdermin and reporter mRNAs into liposomes.

We first assessed the transfection efficiency using EGFP mRNA as a reporter and found comparable uptake and expression between WT and Rabgef1-deficient macrophages (Figure S5B and S5C). We next transfected WT and Rabgef1-deficient macrophages with mRNA encoding both FL and NT gasdermins to evaluate pore forming activities. As expected, transfection of cells with NT gasdermins induced membrane permeability, as assessed by PI positive single cell frequency, compared to the inactive FL gasdermins (Figure S5D-H). Notably, regardless of the gasdermin examined, Rabgef1-deficient macrophages were defective for NT gasdermin-dependent plasma membrane permeability, as compared with WT macrophages. Rabgef1 is therefore a general regulator of the gasdermin family (Figure S5D-H).

Our use of mRNAs to transfect cells with gasdermins offered a simple platform to assess the function of cholesterol in regulating pyroptosis in diverse cellular settings. Given the growing interest in harnessing inflammatory cell death as a potential antitumor strategy, we determined if the link between cholesterol abundance and pyroptosis extends to cancer cells. Similar to our findings in macrophages, MCD treatment of murine B16F10 melanoma cells depleted cholesterol (Figure 6A). MCD treatment led to an increase in membrane permeability across a range of doses of mRNA encoding human gasdermin NTs, with limited increases in membrane permeability for inactive human gasdermin FL variants (Figure 6B-6F). While this trend of cholesterol depletion by MCD sensitizing B16F10 melanoma cancer cells to the active pore forming domains of gasdermins was significant for NT-GSDMB, NT-GSDMC and NT-GSDMD with limited enhancement of membrane permeability for the inactive human FL-GSDMB, FL-GSDMC, and FL-GSDMD (Figure 6C-E), the enhancement of gasdermin pore activity by MCD cholesterol depletion was most apparent in B16F10 melanoma cancer cells treated with human NT-GSDMA and NT-GSDME (Figure 6B and 6F). Our results demonstrate that NT-GSDMA and NT-GSDME have limited intrinsic plasma membrane pore activity by PI staining across a range of mRNA doses that are dramatically increased by MCD treatment. This MCD enhancement of PI staining does not occur in cells transfected with inactive human FL-GSDMA and FL-GSDME (Figure 6B and 6F). Thus, cholesterol dependence of gasdermin pore forming activities generalizes to other cell types including cancer lines and to the pore forming domains of all human gasdermin family members known to form membrane pores (GSDMA, GSDMB, GSDMC, GSDMD, and GSDME).

**Figure 6.**
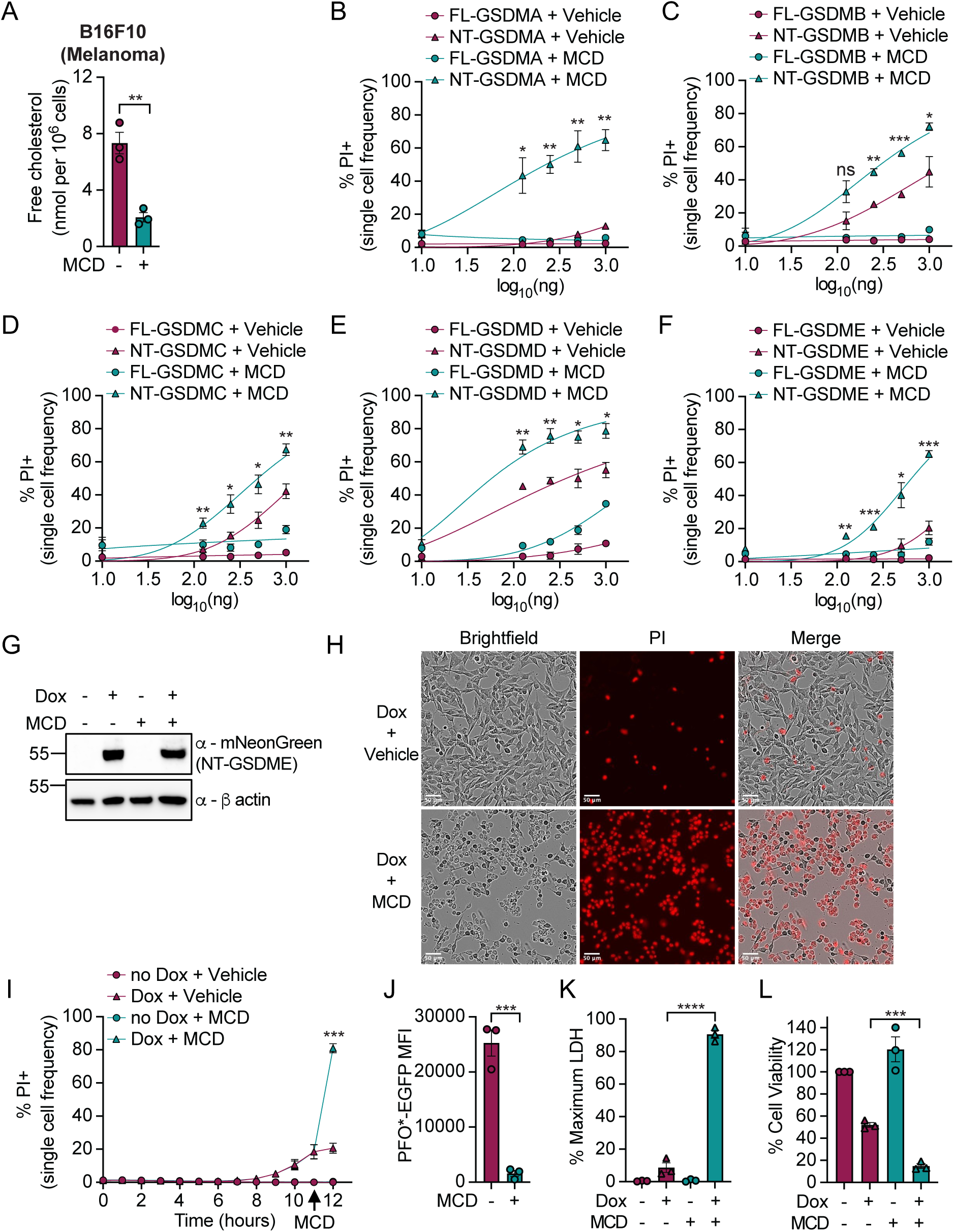
Cholesterol depletion sensitizes cancer cells to pyroptosis by diverse members of the gasdermin family. (A) Free cholesterol quantification in B16F10 melanoma cells treated with or without MCD (7.5 mM) for 1 h. (B-F) PI positive single cell frequency of B16F10 melanoma cells transfected with serial doses of mRNAs encoding FL or NT domains of GSDMA (B), GSDMB (C), GSDMC (D), GSDMD (E), and GSDME (F), together with EGFP mRNA as carrier for 7h, followed with or without MCD (7.5 mM) treatment for 1 h. (G) Western blot analysis of murine NT-GSDME expression in Dox inducible B16F10-NT-GSDME-mNeonGreen cells with or without Dox induction for 11 h, followed with or without MCD (7.5 mM) treatment for 1 h. (H) Live cell images of Dox inducible B16F10-NT-GSDME-mNeonGreen cells labelled with PI after Dox induction for 11 h, followed with or without MCD (7.5 mM) treatment for 1 h. (I) Kinetics of PI positive single cell frequency in Dox inducible B16F10-NT-GSDME-mNeonGreen cells with or without Dox induction for 11 h, followed with or without MCD (7.5 mM) treatment for 1 h. (J) MFI of PFO*-EGFP in Dox inducible B16F10-NT-GSDME-mNeonGreen cells treated with or without MCD (7.5 mM) for 1 h, followed by PFO*-EGFP staining. (K and L) Bulk LDH release (K) and cell viability (L) in Dox inducible B16F10-NT-GSDME-mNeonGreen cells induced with or without Dox for 11 h, followed with or without MCD (7.5 mM) treatment for 1 h. Data are mean ± SEM of three repeats. Immunoblots show representative result of three repeats. Statistical significance was determined by unpaired two-tailed t-test: *p < 0.05; **p < 0.01; ***p < 0.001; ****p < 0.0001.

GSDME was recently shown to potentiate antitumor immune responses^18–20^. To determine if cholesterol depletion sensitizes GSDME-mediated pyroptosis across diverse cancer types, we transfected a panel of murine cancer lines of different tissue sites including pancreatic cancer cells (PDAC) (Figure S6A-D), Lewis lung cancer cells (LLC1) (Figure S6E-H), bladder cancer cells (MB49) (Figure S6I-L), and breast cancer cells (EO771) (Figure S6M-P) with increasing doses of human NT-GSDME mRNA with or without MCD treatment. Consistent with observations in macrophages and B16F10 cells, MCD treatment reduced cellular free cholesterol content (Figure S6A, S6E, S6I and S6M) and also depletes plasma membrane cholesterol (Figure S6B, S6F, S6J and S6N) in diverse cancer cell lines. Transfection with NT-GSDME alone induced limited pore formation, whereas cholesterol depletion enhanced GSDME-dependent plasma membrane permeability (Figure S6C, S6G, S6K and S6O) without affecting GSDME protein production (Figure S6D, S6H, S6L and S6P). These findings suggest that the potentiation of GSDME-mediated pyroptosis by cholesterol depleting agents is a broadly applicable regimen across cancer types.

Based on the apparent inherent resistance of cancer cells to the pore-forming activity of GSDME and the conversion to pyroptotic cell death upon cholesterol depletion, we created a new Dox-inducible cellular model of gasdermin pore formation using the N terminal pore forming fragment of murine GSDME fused to the fluorescent protein mNeonGreen (NT-GSDME-mNeonGreen) in B16F10 melanoma cells expressing Tet3G. This model induces expression of NT-GSDME-mNeonGreen in a Dox-dependent manner (Figure 6G). Similar to mRNA-liposome transfections with human gasdermin variants, B16F10 melanoma cells exhibited delayed responsiveness to NT-GSDME-mediated plasma membrane permeability with minimal PI staining within the first 12 hours of induction (Figure 6H and 6I). Strikingly, MCD treatment of engineered B16F10 melanoma cells in the last hour of NT-GSDME Dox induction causes increased gasdermin-dependent plasma membrane permeability (Figure 6H and 6I). MCD specifically depleted plasma membrane cholesterol, as detected by PFO*-EGFP staining (Figure 6J). Furthermore, MCD treatment of NT-GSDME expressing B16F10 melanoma cells increased pyroptotic lysis (Figure 6K) and reduced cellular viability (Figure 6L) without affecting GSDME abundance (Figure 6G). The process of GSDME-pyroptosis is therefore potentiated by cholesterol depletion.

### Reducing cholesterol concentration enhances NT-GSDME mediated pyroptosis in organoids and tumors

Our results demonstrate that MCD depletion sensitizes cancer cells to GSDME-mediated pyroptosis through mRNA transfection or chemogenetic induction. To validate these findings in a cancer tissue model, we engineered murine *Kras^G12D/+^Trp53^fl/fl^*(KP) pancreatic ductal adenocarcinoma (PDAC) organoids to express Tet3G and TRE3G-promoter upstream of murine NT-GSDME fused to a C terminal mNeonGreen. Engineered KP PDAC organoids were pre-treated with either AY9944 or 25HC, followed by Dox induction of NT-GSDME expression. Dox treatment induced NT-GSDME-mNeonGreen expression in engineered KP PDAC organoids which was not affected by AY9944 or 25HC pre-treatment (Figure 7A and S7A). NT-GSDME expression alone results in minimal plasma membrane permeability seen by PI staining (Figure 7B, 7C, S7B and S7C, Video S4). AY9944 or 25HC pre-treatment decreased plasma membrane cholesterol in KP PDAC organoids (Figure 7D and S7D, Video S5 and S6). Cholesterol depletion by AY9944 or 25HC treatment sensitizes KP PDAC organoids to NT-GSDME-mediated pyroptosis based on increased plasma membrane permeability by PI staining (Figure 7B, 7C, S7B and S7C), increased cellular lysis by LDH release (Figure 7E and S7E), and reduced cell viability by normalized ATP quantification (Figure 7F and S7F).

**Figure 7.**
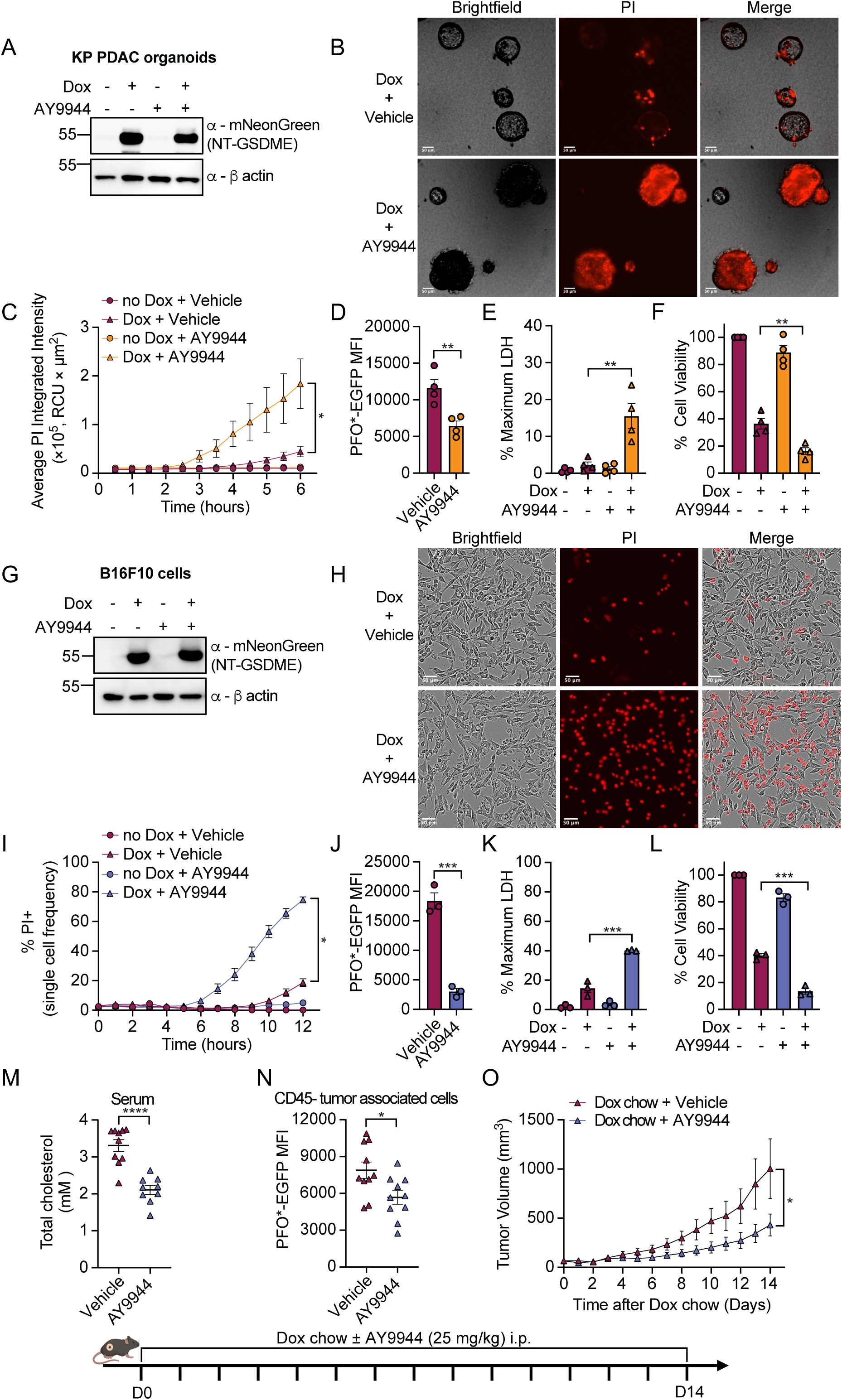
Cholesterol depletion synergizes with GSDME-mediated pyroptosis in cancer organoids and tumor models. (A) Western blot analysis of murine NT-GSDME expression in Dox inducible KP pancreatic organoids pre-treated with or without AY9944 (10 µM) for 4 h, followed with or without Dox induction for 6 h. (B) Live cell images of Dox inducible KP pancreatic organoids labelled with PI and pre-treated with or without AY9944 (10 µM) for 4 h, followed with Dox induction of NT-GSDME expression for 6 h. (C) Kinetics of average PI integrated intensity in Dox inducible KP pancreatic organoids pre-treated with or without AY9944 (10 µM) for 4 h, followed with or without Dox induction for 6 h. (D) MFI of PFO*-EGFP in Dox inducible KP pancreatic organoids treated with or without AY9944 (10 µM) for 10 h. (E and F) Bulk LDH release (E) and cell viability (F) in Dox inducible KP pancreatic organoids pre-treated with or without AY9944 (10 µM) for 4 h, followed with or without Dox induction of NT-GSDME expression for 6 h. (G) Western blot analysis of murine NT-GSDME expression in Dox inducible B16F10-NT-GSDME-mNeonGreen cells pre-treated with AY9944 (10 µM) for 8 h, followed with or without Dox induction of murine NT-GSDME expression for 12 h. (H) Live cell images of Dox inducible B16F10-NT-GSDME-mNeonGreen cells labelled with PI and pre-treated with AY9944 (10 µM) for 8 h, followed with Dox induction of NT-GSDME expression for 12 h. (I) Kinetics of PI positive single cell frequency in Dox inducible B16F10-NT-GSDME-mNeonGreen cells pre-treated with AY9944 (10 µM) for 8 h, followed with or without Dox induction of NT-GSDME expression for 12h. (J) MFI of PFO*-EGFP in Dox inducible B16F10-NT-GSDME-mNeonGreen cells treated with or without AY9944 (10 µM) for 20 h. (K and L) Bulk LDH release (K) and cell viability (L) in Dox inducible B16F10-NT-GSDME-mNeonGreen cells pre-treated with or without AY9944 (10 µM) for 8 h, followed with or without Dox induction of NT-GSDME expression for 12 h. (M) Melanoma bearing C57BL/6J mice were fed Dox chow and treated daily with AY9944 (25 mg/kg, i.p.) or vehicle control for 14 days. Total cholesterol concentration in serum was quantified at the experimental endpoint (n=10). (N) Melanoma bearing C57BL/6J mice were treated daily with AY9944 (25 mg/kg, i.p.) or vehicle control for 14 days. Tumors were collected for PFO*-EGFP staining. MFI of PFO*-EGFP from CD45 negative tumor associated cells were analyzed (n=10). (O) Tumor growth in melanoma bearing C57BL/6J mice fed Dox chow and treated daily with AY9944 (25 mg/kg, i.p.) or vehicle control over time (n=10). For C-F, data are mean ± SEM of four repeats. For I-L, data are mean ± SEM of three repeats. Immunoblots show representative result of at least three repeats. For M-O, data are mean ± SEM and are representative of two repeats. Statistical significance was determined by unpaired two-tailed t-test: *p < 0.05; **p < 0.01; ***p < 0.001; ****p < 0.0001.

To determine if GSDME-mediated pyroptosis was similarly responsive to small molecule modulation of cholesterol metabolism in B16F10 cells, we pre-treated B16F10 cells with either AY9944 or 25HC followed by Dox induction of NT-GSDME expression. Dox treatment induced NT-GSDME-mNeonGreen expression in engineered B16F10 melanoma cells which was not affected by AY9944 or 25HC pre-treatment (Figure 7G and S7G). NT-GSDME expression alone in engineered B16F10 melanoma cells results in minimal plasma membrane permeability seen by PI staining (Figure 7H, 7I, S7H and S7I, Video S7). AY9944 or 25HC pre-treatment decreases plasma membrane cholesterol in B16F10 melanoma cells (Figure 7J and S7J).

Cholesterol depletion by AY9944 or 25HC treatment sensitizes B16F10 melanoma cells to NT-GSDME-mediated pyroptosis based on increased plasma membrane permeability by PI staining (Figure 7H, 7I, S7H and S7I, Video S8 and S9), increased cellular lysis by LDH release (Figure 7K and S7K), and reduced cell viability by normalized ATP quantification (Figure 7L and S7L).

To evaluate whether AY9944 cholesterol depletion synergizes with NT-GSDME to enhance antitumor response *in vivo*, Dox inducible NT-GSDME expressing B16F10 melanoma cells were implanted subcutaneously in the flank of mice. Palpable tumors were allowed to grow to around 50-100 mm^3^ prior to start of treatment. Tumor bearing mice were switched to Dox chow and injected intraperitoneally with vehicle or AY9944 daily for 14 days. AY9944 injection effectively decreased total cholesterol and free cholesterol levels in mouse serum (Figure 7M and S7M). Moreover, AY9944 injection resulted in reduced PM cholesterol in CD45 negative tumor associated cells by PFO*-EGFP staining of homogenized tumors (Figure 7N). Dox induction of NT-GSDME alone demonstrates robust growth of B16F10 melanoma tumors based on raw tumor volume or normalized to tumor volume at the start of treatment (Figure 7O and S7N). The combination of Dox induced tumor cell NT-GSDME expression and cholesterol depletion via AY9944 injection significantly diminishes tumor growth (Figure 7O and S7N) with limited effects on mouse body weight (Figure S7O). These data support a synergistic effect of NT-GSDME production and cholesterol depletion in mediating an antitumor effect *in vivo*.

FL-GSDME is naturally activated by apoptotic signaling specifically via caspase-3 dependent cleavage^18–20^. Caspase-3 signaling is activated by Raptinal or chemotherapy drugs such as Cisplatin^59^. To determine whether cholesterol levels affect natural GSDME-mediated pyroptosis downstream of caspase-3 signaling, we generated stable B16F10 melanoma cell lines expressing mNeonGreen as a control or murine FL-GSDME-mNeonGreen. We pre-treated B16F10-mNeonGreen and B16F10-FL-GSDME-mNeonGreen cells with AY9944 or 25HC followed by Raptinal or Cisplatin treatment to activate caspase-3 and induce GSDME cleavage. Consistent with previous reports, both Raptinal and Cisplatin treatment induces FL-GSDME cleavage (Figure S8A, S8B, S8I and S8J). In B16F10-FL-GSDME cells, cleavage of FL-GSDME results in pyroptotic cell death characterized with cell membrane permeability indicated by PI positive single cell frequency, cell membrane rupture indicated by LDH release, and reduced cell viability (Figure S8C-S8H, S8K-S8P). Notably, reducing cholesterol content by AY9944 or 25HC treatment further enhances FL-GSDME cleavage mediated pyroptosis in B16F10 cells with increased membrane permeability, LDH release, and reduced cell viability (Figure S8C-S8H, S8K-S8P).

These data establish cholesterol as a broad regulator of gasdermin pore activity. Our results also position strategies to produce active gasdermins within target cells, such as our synthetic killer mRNAs or conventional chemotherapies, combined with cholesterol metabolic sensitization as a proof-of-concept strategy for innate immune-based cell death therapy.

## Discussion

Pyroptosis is an important cell death pathway for control of microbial invasion, contextual instruction of adaptive immunity, and pathologic issues in chronic diseases. Much of the prior literature has focused on regulation of pyroptosis from the perspective of transcriptional priming, pathway triggers, and protease-mediated regulation of gasdermin family members to liberate pore forming NT domains from their autoinhibitory CT domains. Recent work from our group and others have begun to uncover post-cleavage regulatory events that dictate pore formation by the gasdermin family of pyroptotic executioners. These include cell biological mechanisms of degradation and repair^60–62^, metabolic and biochemical mechanisms of post-translationa modifications^38,39,63–65^, and positive associations of NT domains with acidic, negatively charged phospholipid species^10,13^.

Herein, we establish membrane cholesterol as an endogenous checkpoint of gasdermin pore formation. While this possibility was previously made plausible biophysically in model membranes with a single gasdermin family member, our current data demonstrate that this principle holds in living cells, including immune cells, cancer cells, organoids, and tumors.

Furthermore, our data suggest that cells can respond to perturbations to modulate cholesterol concentration, with profound impacts on a cell type’s or cell state’s resistance or sensitivity to gasdermin-mediated pyroptosis. We determine that endosome adaptors positively regulate pyroptosis through their role in dictating the content of host plasma membranes, specifically as it pertains to cholesterol. We find endosome adaptor deficiency, such as Rabgef1 knock out macrophages, leads to the accumulation of cholesterol within plasma membranes. This accumulation directly impinges on the ability of NT-GSDMD and other gasdermin NT fragments to oligomerize and form functional pores at the plasma membrane of host cells. Moreover, cholesterol depletion sensitizes resistant cell types such as Rabgef1-deficient macrophages and diverse cancer cell models to pyroptosis by gasdermin NT fragments. While extensive work has uncovered diverse regulatory mechanisms that dictate pyroptotic pathway activation, our study positions cholesterol metabolism as a target to promote pyroptosis in contexts such as cancer therapy. These findings identify cholesterol as the first lipid species shown to restrain pore formation by the pore competent members of the mammalian gasdermin family. Previous studies emphasize the positive determinants of gasdermin targeting and activity, and our results now add a complementary principle to the rules of gasdermin pore formation, namely that membrane cholesterol actively constrains gasdermin executioner function in living immune cells, cancer cells, organoids, and tumors.

Several molecular models are compatible with our data. The most direct interpretation is that plasma membrane cholesterol content alters the physical state of the bilayer, including order and rigidity, thereby raising the barrier to the insertion and oligomerization stages of gasdermin pore formation. Indeed, our cellular biochemistry is consistent with this interpretation. Bulk membrane binding of NT-GSDMD remains intact even in cells with elevated cholesterol, whereas downstream functional aspects of gasdermin pore formation, including oligomerization, membrane permeability, energetic death, and cellular lysis, are inhibited. This represents an important advance in the interpretation of an earlier biophysical report on cholesterol and recombinant GSDMD, in which their assay could not distinguish among binding, insertion, or oligomerization mechanisms; if any of those were inhibited, AFM pore-area graphs would appear similar to their reported results^26^. Across the graded, physiological range of sterol concentrations that cells can occupy, our fractionation data place this cholesterol-sensitive step of gasdermin pore formation downstream of membrane recruitment. Therefore, a reconstituted biophysical system and a cellular *in vivo* system are converging on a similar conclusion from opposite directions, with our cellular system resolving which step of pore formation is controlled by sterol content. This model suggests that membrane cholesterol serves as an intrinsic checkpoint governing whether gasdermin cleavage and upstream death-pathway signaling culminate in inflammatory cell lysis.

Our current model that cholesterol impinges on the efficiency of gasdermin pore formation predicts that the sterol gradient across cellular membranes should dictate where and when gasdermins act within dying cells. In support of this interpretation, organelle membranes with a relative paucity of cholesterol yet containing sufficient acidic phospholipid for gasdermin binding would be expected to be most efficiently targeted, and the plasma membrane, which is often considered the most cholesterol-rich membrane of the cell, should be somewhat refractory. The emerging data on subcellular gasdermin targeting fit this prediction. Indeed, mitochondrial permeabilization by NT gasdermin fragments precedes and accelerates perforation of the plasma membrane^25^. Furthermore, GSDME in neurons, a cell type whose plasma membrane carries unusually high cholesterol, displays a marked preference for mitochondria over the plasma membrane^66^. We propose that the same checkpoint will dictate the endolysosomal and mitochondrial targeting efficiencies now being described for individual gasdermin family members, and measuring sterol content membrane by membrane within a given cell will predict the order in which those membranes fail in response to gasdermins. The corollary at the scale of whole cells is that the cell type, cell state, and cancer lineage each tune gasdermin output, in part, through their sterol and lipid trafficking programs. We find this directly in malignant cells, where particular models highlighted here with elevated cholesterol resist GSDME pyroptosis despite producing abundant cleaved executioner. A checkpoint of this kind may reconcile a portion of the otherwise puzzling variation in how readily different cells die by pyroptosis for similar amounts of active gasdermin.

Our findings place endosomal trafficking at the center of metabolic control over inflammatory cell death. Whether these same adaptors and cholesterol mechanisms also dictate the activity of other membrane-disrupting agents, such as the necroptotic executioner MLKL, the pan-lysis executioner NINJ1, and the recently described pore-forming family of APOLs, awaits further study. Certainly, cholesterol control of membrane permeability by membrane-disrupting proteins is not absolute, as cholesterol content positively regulates pore formation by host perforin pores downstream of cytolytic lymphocytes, including T cells and NK cells, similar to bacterial cholesterol-dependent cytolysins^67,68^.

Our identification of cholesterol as a gasdermin suppressor carries direct consequences for cancer biology, as we have begun to explore in this study. Many tumors remodel cholesterol synthesis and uptake, and the selective advantages usually invoked are cell-intrinsic proliferation, invasive capacity, and cell-extrinsic suppression of local immunity. We suggest an additional one, which is resistance to lytic death executioners of the gasdermin family. Our data directly demonstrate that the metabolic state of a cancer cell membrane confers protection against GSDME-mediated pyroptosis even when apoptotic caspases are activated and pore-competent GSDME fragments are robustly produced. Importantly, cholesterol-depleting strategies convert otherwise resistant cancer cells, organoids, and tumors into pyroptotic targets. Read as a stratification problem, this predicts three clinically distinguishable groups.

First, tumors without appreciable gasdermin expression are not expected to respond well to pyroptosis-activating therapies. Second, tumors with high gasdermin expression and low cholesterol biosynthetic output should be intrinsically favorable substrates for pyroptosis-activating therapy and may respond without metabolic intervention. Third, tumors with high gasdermin expression and high cholesterol biosynthetic signatures should require metabolic sensitization before a pyroptotic trigger can be converted into immunogenic lysis. Tumors that combine high gasdermin expression with high cholesterol signatures are the group in which we would expect the clearest benefit from adding approved cholesterol-lowering agents as a rational combination approach. Statins and the newer PCSK9 inhibitors^69^ are the obvious candidates for these combination trials, and the tolerability of both is well established, permitting testing alongside cytotoxic and precision oncogene inhibitor regimens. These observations add to the pro-cancer and anti-immunity functions of tumor-associated metabolic reprogramming.

Cholesterol content in cancer cells and tumors is currently purported to support growth, invasion, and immune evasion, and our data now add a role of membrane cholesterol in active suppression of inflammatory forms of cell death.

This same logic likely extends to immunotherapy. GSDME and GSDMB are activated within target cancer cells by granzymes delivered from cytotoxic lymphocytes^20,21^, which places killer immune cell engagement directly upstream of a step that membrane cholesterol controls. Lowering cancer cell cholesterol would therefore be expected to raise the fraction of lymphocyte engagements that end in pyroptotic lysis rather than in non-lytic or apoptotic outcomes. Thus, cholesterol-lowering therapies may synergize with cancer therapies that aim to activate killer immune cells against target cancer cells, including checkpoint blockade, adoptive tumor infiltrating lymphocyte transfer, CAR T cells, and personalized cancer vaccines. One caveat to this prediction requires consideration and empirical testing. Perforin itself has been suggested to depend on target membrane cholesterol for the initial monomer binding stage of pore formation, so the potential that a sufficiently aggressive reduction in membrane cholesterol might, in principle, blunt the delivery step on which the granzyme entry relies. Whether a therapeutic window separates these two scenarios is an empirical question that should be answered to enable more complex combinations, including cholesterol-depleting agents, conventional cancer therapies that can activate gasdermins, and immunotherapies.

Taken together, this work establishes cholesterol metabolism and membrane cholesterol as intrinsic checkpoints for membrane attack by gasdermin pores. The integration of metabolic state, membrane trafficking, and protease activity allows cells to calibrate the intensity and ultimate outcome of inflammatory death programs. From a therapeutic standpoint, our findings identify metabolic sensitization as a requirement for fully exploiting pyroptosis-activating strategies in cancer, vaccination, and infection. Cholesterol biosynthetic signatures and statin use may prove useful for stratifying patient cohorts alongside oncogene driver status, tumor suppressor status, BH3 profiling, and the current biomarkers that are used to predict responsiveness to chemotherapy, immunotherapy, precision inhibitors, and personalized vaccines. More broadly, these data argue cholesterol metabolism and membrane content are levers through which inflammatory fate decisions are controlled in immunity and cancer.

## Materials and Methods

### KEY RESOUCES TABLE

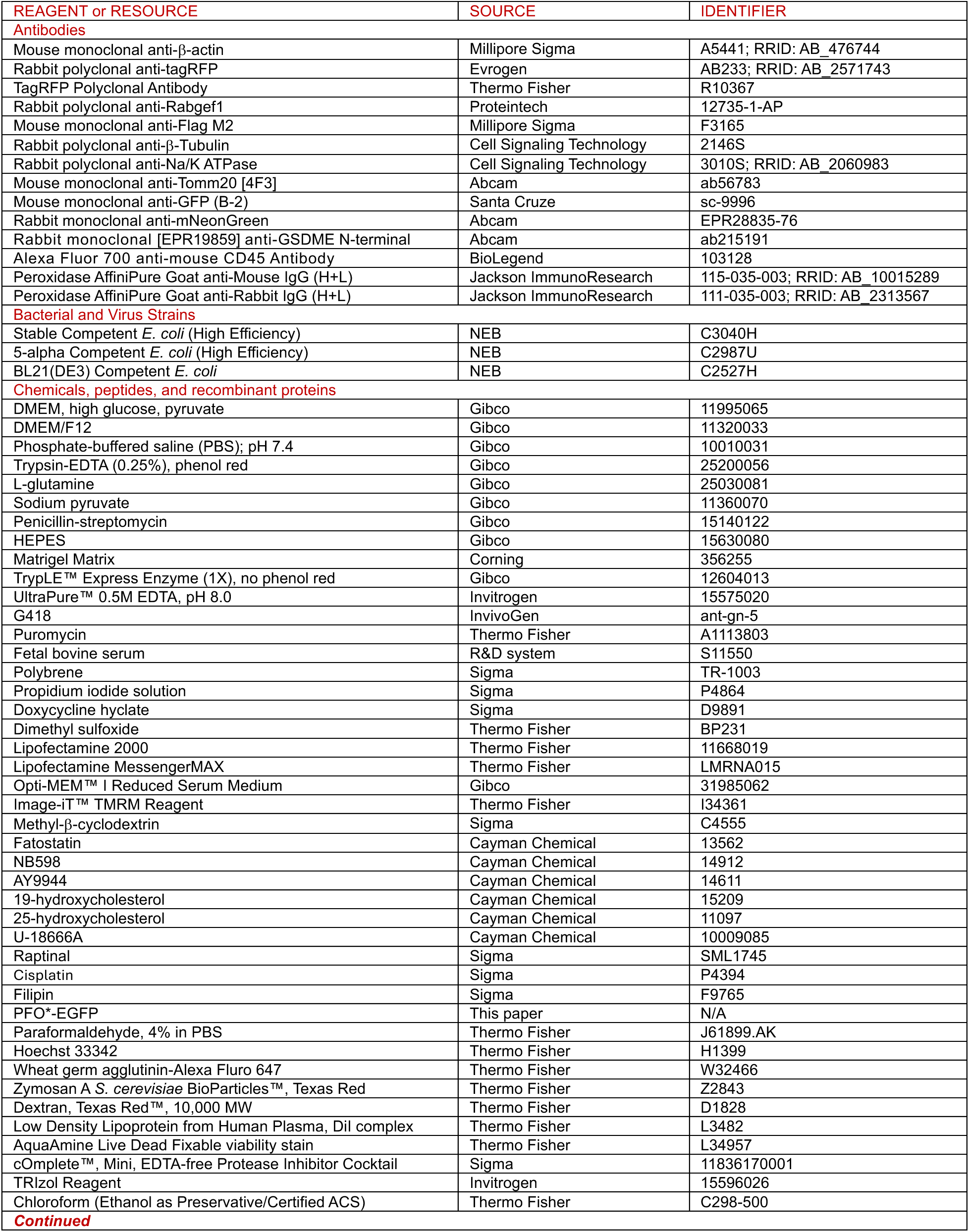
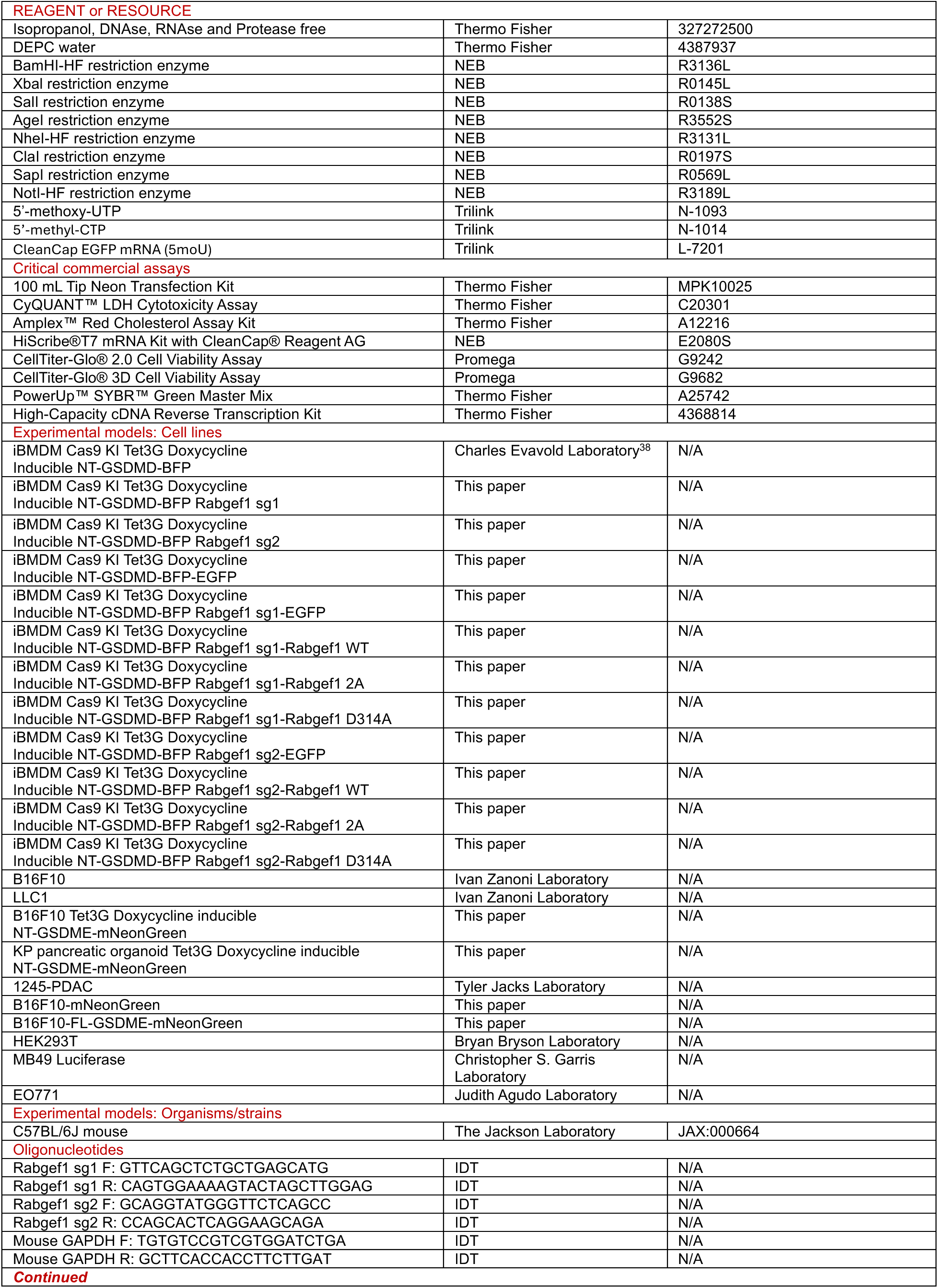
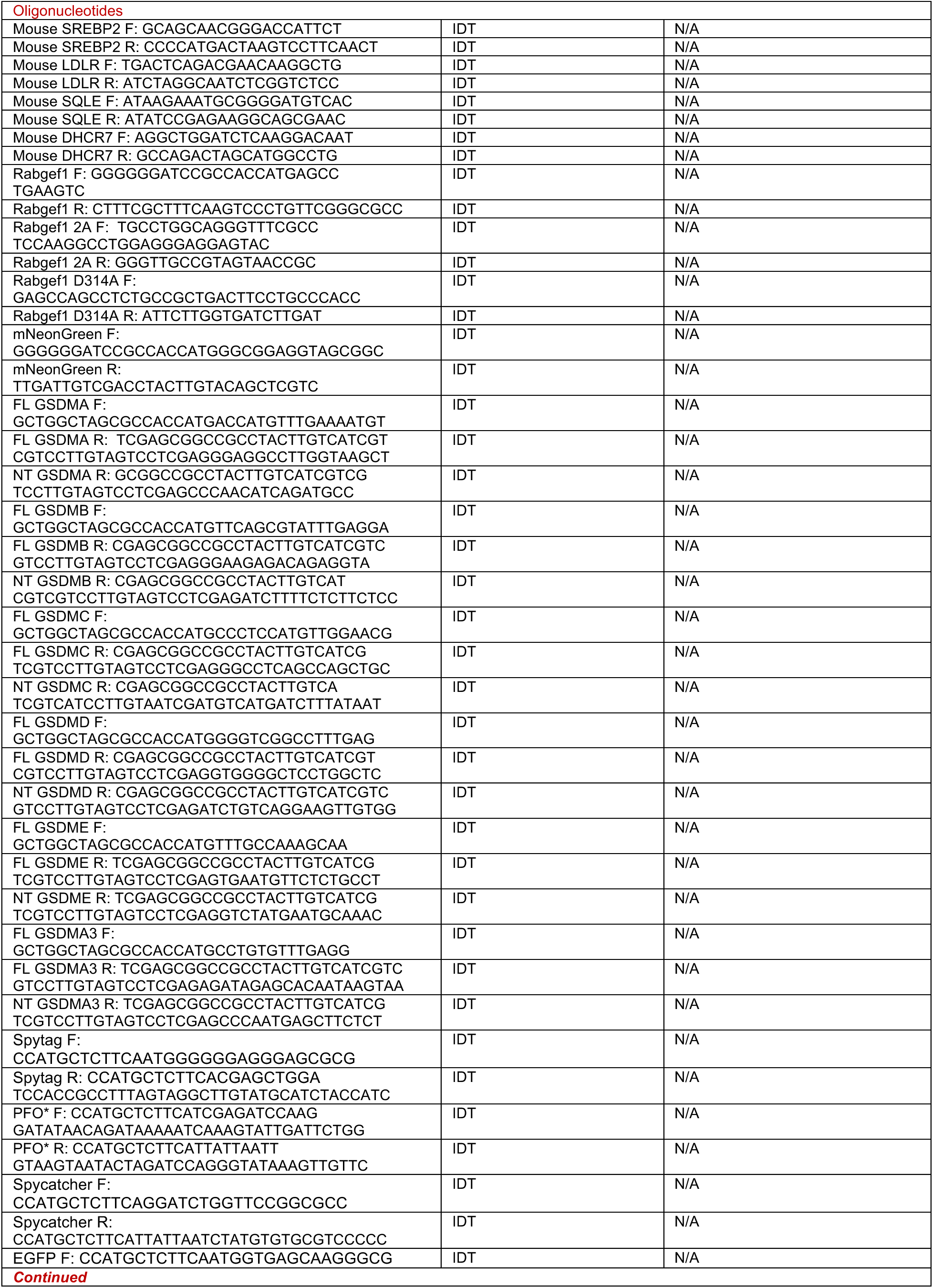
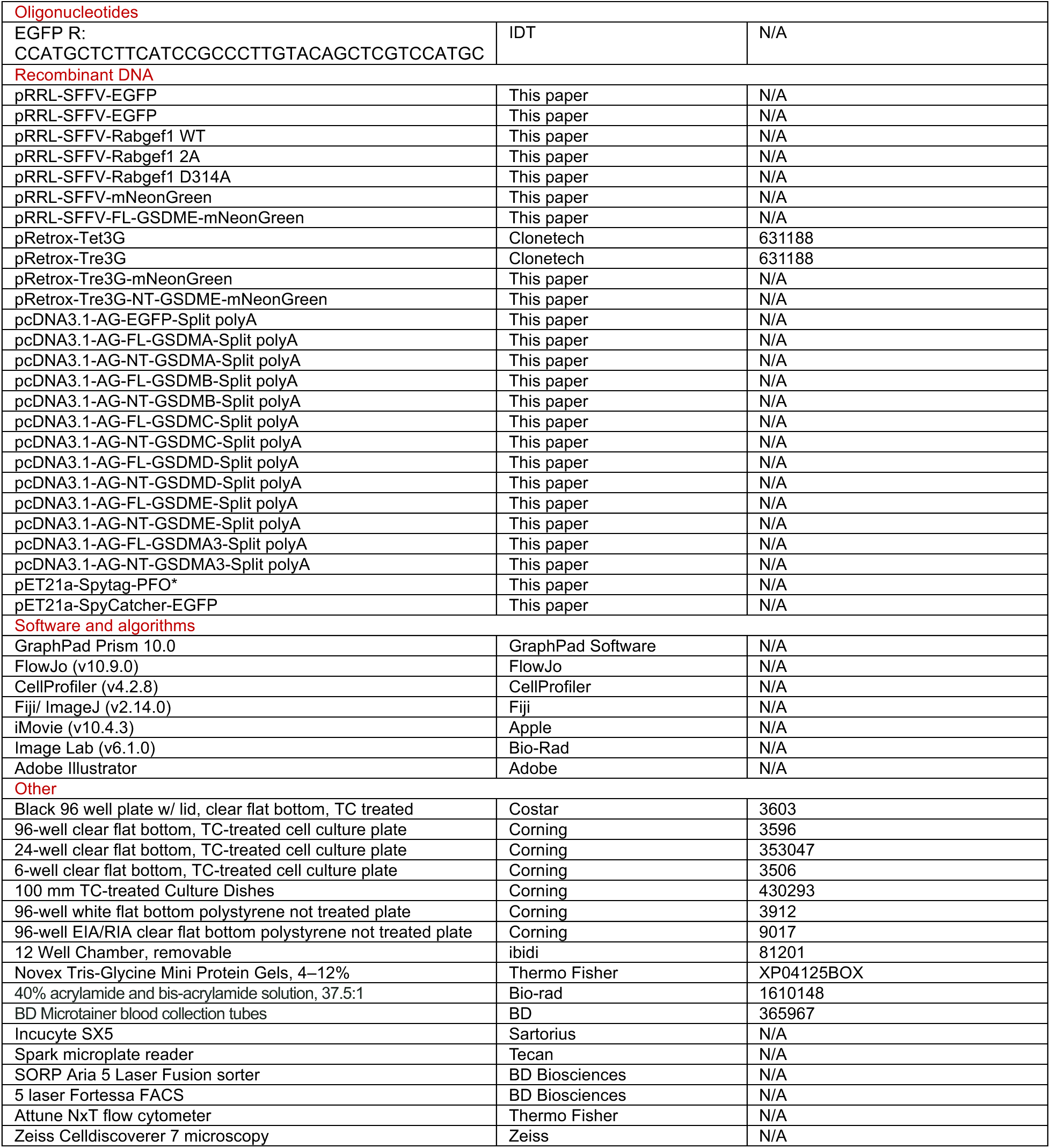

## EXPERIMENTAL MODEL AND SUBJECT DETAILS

### Cell culture and KP pancreatic organoids generation

All cells were cultured in humidified incubators containing 5% CO_2_ at 37°C. Immortalized bone marrow-derived macrophages (iBMDMs), HEK293T, B16F10, 1245-PDAC, LLC1 and MB49 cells were maintained in DMEM supplemented with 10% fetal bovine serum (FBS), 2 mM L-glutamine, 100 U/mL penicillin-streptomycin and 1 mM sodium pyruvate. EO771 cells were maintained in DMEM supplemented with 20 mM HEPES, 10% FBS, 100 U/mL penicillin-streptomycin and 2 mM L-glutamine.

KP pancreatic organoids were generated as previously described^70^. Briefly, the pancreas from a C57BL/6J Kras^LSL-G12D/+^ Trp53^flox/flox^ (KP) mouse was finely minced and digested with 125 U/mL collagenase IV in HBSS for 20 min at 37°C. Digested tissue was filtered through a 70 μm filter and washed three times with PBS by centrifuging at 850 x g for 3 min. The resulting cell pellet was resuspended in 100% growth-factor reduced Matrigel matrix and solidified at 37°C before addition of pancreatic organoid media, which permits the selection of ductal cells over several passages. Organoids were transformed by Cre recombination by resuspending cells in 85% Matrigel matrix spiked with 15% Ad5-CMV-Cre virus diluted in PBS at an MOI of 500. Recombined cells were selected with addition of 10 μM Nutlin-3a (Sigma-Adlrich) to promote apoptosis of remaining cells expressing p53. KP pancreatic organoids were maintained in a base of advanced DMEM/F-12 medium containing 2 mM GlutaMAX, 100 U/mL penicillin-streptomycin, 20 mM HEPES, and further supplemented with pancreatic organoid media containing 0.5 μM A83-01, 0.05 μg/mL mEGF, 0.1 μg/mL hFGF-10, 0.01 μM hGastrin I, 0.1 μg/mL mNoggin, 1.25 mM N-acetylcysteine, 10 mM Nicotinamide, R-spondin 1-conditioned medium, B27 supplement and 10.5 μM Y-27632. To passage organoids, domes were mechanically disrupted and centrifuged at 850 x g for 3 min. Supernatant was carefully aspirated while leaving the layer of matrigel, which was then digested using TrypLE Express Enzyme for at least 15 min at 37°C. Cells were centrifuged at 850 x g for 3 min and resuspended in a solution of 85% Matrigel matrix/15% advanced DMEM/F12 medium. Domes were allowed to solidify at 37°C for at least 10 min before addition of pancreatic organoid media.

### Reagents and antibodies

Propidium iodide solution (1 mg/mL) was used at a final concentration of 3.3 µg/mL (1:300 dilution). Image-iT™ TMRM reagent (100 µM) was used at a final concentration of 150 nM (1:666 dilution). Doxycycline hyclate was dissolved in sterile water at a concentration of 2 mg/mL and used at a final concentration of 1 µg/mL (1:2000 dilution). Methyl-β-cyclodextrin (MCD) was dissolved in sterile water at a concentration of 200 mM and used at a final concentration of 10 mM for iBMDMs or 7.5 mM for cancer cells. Fatostatin was dissolved in DMSO at a concentration of 20 mM and used at a final concentration of 20 µM (1:1000 dilution). NB598 was dissolved in DMSO at a concentration of 10 mM and used at a final concentration of 2.5 µM (1:4000 dilution). AY9944 was dissolved in DMSO at a concentration of 10 mM and used at a final concentration of 10 µM (1:1000 dilution). 19-hydroxycholesetrol (19HC) and 25-hydroxychoelsterol (25HC) were dissolved in ethanol at a concentration of 10 mM and used at a final concentration of 10 µM (1:1000). U18666A was dissolved in DMSO at a concentration of 10 mM and used at a final concentration of 2 µM (1:5000 dilution). Raptinal was dissolved in DMSO at a concentration of 10 mM and used at a final concentration of 5 µM (1:2000 dilution). Cisplatin was dissolved in PBS at a concentration of 10 mM and used at a final concentration of 160 µM (1:62.5 dilution).

The following antibodies were used as primary antibodies for immunoblotting: mouse monoclonal anti-β-actin (Sigma, 1:2000), rabbit polyclonal anti-tagRFP (Evrogen, Thermo Fisher, 1:1000), rabbit polyclonal anti-Na/K+ ATPase (Cell Signaling, 1:1000), rabbit polyclonal anti-Tubulin (Cell Signaling, 1:1000), mouse monoclonal anti-Tomm20 (Abcam, 1:1000), mouse monoclonal anti-GFP (Santa Cruz, 1:1000), rabbit polyclonal anti-Rabgef1 (Proteintech, 1:1000), rabbit monoclonal anti-GSDME (Abcam, 1:1000), rabbit polyclonal anti-mNeonGreen (Abcam, 1:1000), mouse monoclonal anti-Flag M2 (Sigma, 1:1000). HRP-conjugated goat anti-mouse and goat anti-rabbit secondary antibodies were diluted by 1:3000.

### Plasmid construction

DNA encoding the murine Rabgef1 isoform C was ordered from Twist Bioscience. Rabgef1 WT construct was cloned into pRRL-SFFV vector containing T2A-EGFP using BamHl and Xbal restriction sites. Point mutations (C35A, C38A and D314A) were introduced using the Q5 mutagenesis kit according to manufacturer’s protocols. The coding sequences of full-length and N terminal fragment of human GSDMA, GSDMB, GSDMC, GSDMD, GSDME and murine GSDMA3 were amplified from the indicated plasmids (provided by Feng Shao’s lab) by PCR and cloned into pcDNA3.1-AG backbone containing modified 5’UTR, 3’UTR and split polyA tail for in vitro transcription using NheI and NotI restriction sites. The coding sequences of mNeonGreen and murine NT-GSDME-mNeonGreen were amplified by PCR and cloned into pRETROX-Tre3G plasmids using ClaI and NotI restriction sites. The coding sequences of mNeonGreen and murine FL-GSDME-mNeonGreen were amplified by PCR and cloned into pRRL-SFFV vector using AgeI and SalI restriction sites. PFO*, EGFP, SpyTag and SpyCatcher were amplified by PCR and cloned into pEG2206 vector using SapI restriction sites. Primers used for plasmids construction were shown in the oligonucleotides section.

All constructs were verified by Sanger sequencing of inserts and / or Sanger sequencing of difficult templates for split polyA tails, and full plasmid sequencing.

### Human Cell-Map BioID Enrichment Analysis

The most recent “significant preys” dataset was downloaded from the Human Cell-Map BioID database (https://cell-map.org/download, May 2025). Next, a pi value was calculated for each screen hit by multiplying the log_2_FC with the -log_10_(P value). The list of hits with a pi value > 2 were tested for significant enrichment or de-enrichment of each CellMap non-negative matrix factorization (NMF) defined subcellular localization as calculated by the original authors^40^.

Enrichment or depletion was tested by a two-sided Fisher exact test in R, and P-values were adjusted using the Benjamini-Hochberg procedure.

### Generation of Rabgef1 KO cell lines

sgRNAs targeting murine Rabgef1 (sg1: CACTCGGTCCGTCTCAATGC; sg2: ACCACCATGTCGGACACTTC) were ordered from IDT and resuspend into nuclease-free water at a final concertation of 100 µM. Rabgef1 KO cells were generated by electroporation of synthetic sgRNAs into Dox inducible Cas9-expressing iBMDMs using Neon transfection system (Thermo Fisher) with the conditions: 1400 V, 10 ms, 2 pulses, as previously described^38^. Briefly, 1.2E6 cells were suspended into 120 µL of R buffer, mixed with 2.4 µL of sgRNA, and 100 µL mixture containing 1E6 cells was used for electroporation with Neon electroporation pipette tip. Cells were then dispensed directly to a tube containing 900 µL pre-warmed complete DMEM and then transferred to a 6-well plate containing 2 mL pre-warmed complete DMEM. Following 2 days culturing, the cells were collected for assessing bulk knockout efficiency by western blot and subcloned by diluting to 3 cells/mL medium to obtain complete knockout cells. The Rabgef1 expression in subcloned cells was identified by immunoblot. The knockout efficiency of subcloned cells was evaluated by Sanger sequencing as described previously^38^. Briefly, genomic DNA was extracted from 1E5 cells using 50 µL of QuickExtract DNA extraction solution (Lucigen) according to manufacturer’s protocol. Around 500 bp genomic regions flanking the Rabgef1 sgRNA target sequence were amplified by PCR for Sanger sequencing and the sequencing traces were analyzed by ICE software^71^. The knockout efficiency of Rabgef1 sg1 iBMDMs are predicted at 100% and Rabgef1 sg2 iBMDMs are predicted at 91%. The primers used for PCR were shown in oligonucleotides section. The frequency of BFP positivity in WT and Rabgef1-deficient macrophages after Dox induction for 8 h was determined on a Fortessa flow cytometer and analyzed using FlowJo 10.9.0. The intensity of NT-GSDMD after Dox induction for 8 h was determined by western blot.

### Lentiviral/Retrovirus transduction

To reconstitute Rabgef1 in Rabgef1-deficient iBMDMs, 2E6 of HEK293T cells were seeded in 10 cm dishes. On the next day, the cells were transfected with pMD2.G (3 µg), psPAX2 (6 µg) and transfer plasmids pRRL-SFFV encoding Rabgef1 variants (9 µg) using Lipofectamine 2000. 6 h after transfection, the medium was changed to 5 mL fresh culture medium. 48 h after transfection, the lentiviral supernatant was collected, spun down at 400 x g for 5 min to remove cell debris and filtered through 0.45 µm PVDF syringe filter. 1E6 of WT and Rabgef1-deficient iBMDMs were transduced with 4 mL lentiviral supernatant containing polybrene at the concentration of 5 µg/mL, followed by spininfection at 1250 x g for 45 min at 37°C. 3 mL fresh culture medium was added to each well followed with spininfection. Change polybrene containing medium to fresh culture media in the next morning. Because EGFP expression correlated with the transgene expression, the EGFP positive cells were sorted twice for control of transgene expression level. Rabgef1 expressing level was confirmed by immunoblot.

To construct mNeonGreen and murine FL-GSDME-mNeonGreen stable expressing B16F10 cells, the same protocol was used as described above to generate lentivirus encoding mNeonGreen and FL-GSDME-mNeonGreen. B16F10 cells were transduced with the lentiviral supernatant via spininfection and mNeonGreen positive cells were sorted.

To construct Dox inducible murine NT-GSDME-mNeonGreen expression in B16F10 cell lines, three rounds of pantropic retroviruses were transduced. First, Tet3G stable expressing B16F10 cells were generated. 2E6 of HEK293T cells were seeded in 10 cm dishes. On the next day, HEK293T cells were transfected with pMD2.G (3 µg), pCL-Eco (6 µg), and pRETROX Tet3G (9 µg) using Lipofectamine 2000. 6 h after transfection, the medium was changed to 5 mL fresh complete medium. 48 h after transfection, the retroviral supernatant was collected, spun down and filtered through 0.45 µm syringe filter. 5E5 of B16F10 cells seeded in 6-well plate one day before were transduced with 4 mL retroviral supernatant containing polybrene at the concentration of 5 µg/mL, followed by spininfection at 1250 x g for 45 min at 37°C. 3 mL fresh culture medium was added to each well followed with spininfection. Fresh culture medium was changed in the next morning. 36-48 h after transduction, the transduced cells were selected by growth in culture medium containing 2 mg/mL G418. Second, Dox inducible murine NT-GSDME-mNeonGreen in Tet3G expressing B16F10 cells were generated. The retrovirus was produced with the same protocol described above in HEK293T cells transfected with pMD2.G (3 µg), pCL-Eco (6 µg) and pRETROX Tre3G encoding murine NT-GSDME-mNeonGreen (9 µg). B16F10 cells stably expressing Tet3G were transduced twice with retroviruses expressing murine NT-GSDME-mNeonGreen. The transduced B16F10 cells were selected by growth in the medium containing 2 mg/mL G418 and 2 µg/mL puromycin. Dox inducible murine NT-GSDME-mNeonGreen expressing B16F10 cells were subcloned by limit dilution to get the subclonal cells.

To construct Dox inducible murine NT-GSDME-mNeonGreen expression in KP pancreatic organoids, KP pancreatic organoids were transduced with three rounds of pantropic retroviruses. First, Tet3G stable expressing KP pancreatic organoids were generated. Retroviruses expressing Tet3G generated as described above was concentrated and added to TrypLE enzyme digested cell pellets with 85% Matrigel matrix before plating domes. Organoids were maintained and passaged with KP pancreatic organoid culturing medium containing 0.4 mg/mL G418. Second, Dox inducible murine NT-GSDME-mNeonGreen in Tet3G expressing KP pancreatic organoids were generated. Two rounds of retroviruses expressing NT-GSDME-mNeonGreen were transduced in Tet3G expressing KP pancreatic organoids. KP pancreatic organoids were maintained with the medium containing 0.4 mg/mL G418 and 1 µg/mL puromycin.

### GSDMD oligomerization assay

The protocol was described previously^38^. Briefly, 1E6 of WT and Rabgef1-deficient macrophages were seeded in 12-well plate. On the next day, the medium was changed to 1 mL of Opti-MEM containing 1 µg /mL Dox to induce NT-GSDMD expression for 8 h. 200 µL 6 x non-reducing SDS loading buffer was added directly to each well for collecting the protein in both supernatant and cell lysate. The samples were split equally into two tubes and 5% BME or water was added to generate reducing and non-reducing blot samples. All samples were boiled at 65°C for 10 min. The NT-GSDMD oligomers were separated by SDS-PAGE on a 4%-12% acrylamide gradient gel and reduced NT-GSDMD monomers were separated by 10% SDS-PAGE gel.

### Subcellular fractionation

The protocol of subcellular fractionation was described previously^38^. Briefly, 10E6 of WT and Rabgef1-deficient macrophages were seeded in 10 cm dishes. On the next day, NT-GSDMD expression was induced in macrophages cultured with medium containing Dox at the concentration of 1 µg /mL for 8 h. Cells were collected with PBS supplemented with 4 mM EDTA and washed with hypotonic buffer containing protease inhibitors (10 mM Tris pH 7.4, 1.5 mM MgCl_2_, 10 mM KCl, complete mini protease inhibitors). The cells were spun down at 400 x g for 5 min at 4°C and pellet was resuspended in 750 µL hypotonic buffer followed by incubation in ice for 30 min. The cells were homogenized by passing 50 times through a 26G needle and the nuclei and cell debris were removed by centrifugation at 2500 x g for 5 min at 4°C. 100 µL of the supernatant was taken as whole cell lysate (WCL). The remaining supernatant was transferred to new tubes and centrifuged at 17000 x g for 30 min at 4°C to separate the cellular membrane from cytosolic fractionation. The supernatant was transferred to new tubes as cytosolic fractionation. The cellular membrane was washed with hypotonic buffer for three times. The input control, cellular membrane, and cytosolic fraction were lysed with 6 x non-reducing SDS loading buffer. The samples were split equally into two tubes and 5% BME or water was added to generate reducing and non-reducing blot samples. All protein samples were heated at 65°C for 10 min. The samples were separated by 10% SDS-PAGE gel. Tubulin and Na^+^/K^+^ ATPase were used as markers of cytosol and membrane fraction.

### Mitochondria isolation

Mitochondria isolation was performed according to the manufacture’s protocol. Briefly, 20E6 of WT and Rabgef1-deficient macrophages were seeded in two 10 cm dishes. On the next day, NT-GSDMD expression was induced with Dox at the concentration of 1 µg/mL for 8 h. Cells were collected with PBS and spun down at 400 x g for 5 min at 4°C. The cell pellet was incubated with 800 µL reagent A for 2 min on ice. 10 µL reagent B was added directly to each tube and incubate on ice for 5 min followed with 800 µL reagent C treatment. The samples were centrifuged at 700 x g for 10 min at 4°C to remove cell debris. 100 µL of supernatant was kept as whole cell lysate (WCL). The remaining supernatant was transferred to new tubes and spun down at 3000 x g for 15 min to get pure mitochondria. The supernatant was transferred to new tubes as cytosol fraction and the mitochondrial pellet was washed with reagent C. The input control, mitochondrial pellet, and cytosol fraction were lysed with 6 x non-reducing SDS loading buffer. The samples were split equally into two tubes and 5% BME or water was added to generate reducing and non-reducing blot samples. The samples were separated by 10% SDS-PAGE gel. Tubulin and Tomm20 were used as markers of cytosol and mitochondria fraction.

### Propidium iodide staining and TMRM staining

To quantify the cell plasma membrane permeability, the cells were stained with PI at the final concentration of 3.3 µg/mL (1:300 dilution). PI positive single cell frequency was determined over time by SX5 Incucyte live cell imaging system and analyzed based on the cell segmentation in brightfield image.

To assess the mitochondria membrane potential as an indicator of mitochondria condition, WT and Rabgef1-deficient macrophages were stained with TMRM at the final concentration of 150 nM for 40 min. After the incubation, the cells were washed twice with PBS and NT-GSDMD expression was induced with Dox for 8 h. The TMRM positive single cell frequency was analyzed with the Incucyte software based on the cell segmentation in brightfield image. The relative TMRM intensity was calculated by TMRM intensity at 8 h of NT-GSDMD induction normalized to 0 h of NT-GSDMD induction.

### LDH release assay

LDH release was quantified using the CyQuant LDH cytotoxicity assay kit from Thermo Fisher following the manufacturer’s protocol. Briefly, 50 µL/well supernatant was transferred to a 96-well plate containing 50 µL LDH assay buffer and incubate the plate at room temperature for 30 min protected from light. The reaction was stopped with 50 µL of stop solution and measured with the absorbance at 490 nm and 680 nm on a Tecan Spark reader. The maximum LDH release was normalized to lysis control wells.

### Cell viability assay

Cell viability as defined by normalized ATP quantitates from lysed wells of macrophages and B16F10 cells was determined by the CellTiter-Glo Luminescent Cell Viability Assay (Promega, G7571) according to the manufacture’s protocol. Briefly, equal volume of CellTiter-Glo reagent was added to cell culture medium and mixed for 2 min on shaker to induce cell lysis. Luminescence was recorded using Tecan Spark plate reader with an integration time of 0.5 second per well.

The viability of KP pancreatic organoids was determined by CellTiter-Glo 3D Cell Viability Assay kit (Promega, G9681) according to the manufacture’s protocol. Briefly, equal volume of CellTiter-Glo reagent was mixed with cell culture medium for 5 min to induce cell lysis and stabilized for additional 25 min. Luminescence was recorded using Tecan Spark plate reader with an integration time of 0.5 second per well.

### Phagocytosis, pinocytosis and receptor-mediated endocytosis assay

To assess phagocytosis, the indicated cells were incubated with Texas Red conjugated Zymosan A *S. cerevisiae* BioParticles (Invitrogen) at a 1:4000 dilution for 8 h. Phagocytic uptake was monitored over time using SX5 Incucyte live cell imaging system. The frequency of Zymosan-positive cells and fluorescence intensity were quantified using Incucyte analysis software.

To evaluate pinocytosis, the indicated cells were incubated with Texas Red-labeled dextran (10,000 MW, Invitrogen) at a final concentration of 0.5 mg/ mL for 8 h. The Dextran uptake was analyzed by flow cytometry. The percentage of dextran positive cells and MFI were quantified.

To measure receptor-mediated endocytosis, the indicated cells were incubated with Dil labeled LDL (Invitrogen) at a final concentration of 2 µg/ mL for 8 h. The Dil LDL uptake was quantified by flow cytometry. The percentage of Dil LDL positive cells and MFI were determined.

### Cholesterol quantification

Cholesterol concentration of cultured cells was quantified with Amplex Red Cholesterol Assay kit (Invitrogen, A12216) following the manufacturer’s protocol. Briefly, 1E6 cells were seeded in 12 well plate. On the next day, the cells were treated as required. After treatment, the cells were washed three times with PBS and 1E6 cells were collected for cholesterol quantification. The cells were lysed in 500 µL water for 30 min at 37°C before quantification followed by dilution with 2 x reaction buffer. Free cholesterol was measured by incubating 50 μL of the Amplex Red working solution containing 300 μM Amplex Red reagent, 2 U/mL HRP and 2 U/mL cholesterol oxidase with 50 μL samples for 30 min at 37°C. The absorbance was measured using Tecan Spark plate reader with excitation at 560 nm and emission at 590 nm.

Total and free cholesterol concentration in mouse serum was quantified with the same kit described above. Serum was diluted by 1:400 and 1:200 with 1 x reaction buffer respectively for total and free cholesterol quantification. Free cholesterol was quantified with Amplex Red working solution containing 300 μM Amplex Red reagent, 2 U/mL HRP and 2 U/mL cholesterol oxidase and total cholesterol was quantified by adding extra 0.2 U/mL cholesterol esterase in the Amplex Red working solution.

### PFO*-EGFP protein expression, purification, and conjugation

Plasmids encoding SpyTag-PFO* and SpyCatcher-EGFP were transformed into competent *E. coli* BL21. A single colony was inoculated into 66 mL LB medium supplemented with 100 µg/mL carbenicillin and cultured at 225 rpm shaking for overnight at 37°C. On the next morning, 60 mL culture medium was equally inoculated into 10 flasks with 600 mL of 2 x YT broth supplemented with carbenicillin. The stage of bacteria growth was determined with absorbance measurement after 3-5 h culture. When OD600 of bacteria was 0.4-0.8, the transgene expression was induced with 1 mM IPTG at 225 rpm shaking for overnight at 37°C. The bacteria were collected by centrifugation and pellet was resuspended with 240 mL wash buffer (20 mM Tris, 150 mM NaCl, 5 mM imidazole) supplemented with 240 µL of 1.5 mM pepstatin, 24 µL of 10 mM leupeptin, 240 mg of lysozyme and 480 µL of 0.5 M TCEP and incubated on a roller mixer for 1 h at 4°C. The suspension was sonicated with Branson SFX550 sonicator for 2 min (100% amplitude, 10 sec pulses, 1 min between pulses). The lysate was transferred to 50 mL falcon tubes and clarified by centrifugation at 30,000 x g for 30 min at 4°C. The clarified lysate was pooled in a new container, combined with 6 mL of washed cobalt talon resin slurry, and incubated on a roller mixer for 1 h at 4°C. The lysate and resin slurry were transferred to a gravity column and flow through discarded. The resin was washed twice with 30 mL wash buffer supplemented with 1 mM TCEP and eluted twice with 10 mL elution buffer (20 mM Tris, 150 mM NaCl, 150 mM imidazole and 1 mM TCEP, PH 7.4). Eluents were pooled and stored at 4°C for overnight. On the next day, the eluents were centrifuged at 4300 x g for 10 min at 4°C to remove precipitate and the supernatant was transferred to 10 kDa (SpyCatcher-EGFP) or 30 kDa (SpyTag-PFO*) concentrator and centrifuged until 200-450 µL volume remained at 4300 x g for 30-90 min at 4°C. The concentrated eluent was transferred to a 0.2 μm microcentrifuge filter and centrifuged at 16,000 x g for 5 min at 4°C. Filtered concentrated eluent was loaded into a Cytiva AKTA Pure FPLC with an S200 column (Cytiva, 28990944), fraction collector (Cytiva, F9-C), and PBS mobile phase (Corning, 21-031-CV) supplemented with 1 mM TCEP. The FPLC was run at 0.75 mL/min with 23.5 mL buffer and 0.5 mL fractions were collected. After FPLC purification, relevant fractions were pooled, and the concentration was determined with nanodrop. And the protein was verified by SDS-PAGE gel.

To get PFO* protein conjugated with EGFP, SpyTag and SpyCatcher were reacted with the molar ratio=1:4 to ensure that all SpyTag-PFO* was conjugated. SpyTag-PFO* and SpyCatcher-EGFP were mixed by the molar ratio and incubated overnight at 4°C. PFO*-EGFP protein was concentrated through a 30 kDa concentrator to the volume of 200-450 µL, filtered in a microcentrifuge filter tube, and purified via FPLC as described above. The desired fractions were pooled, concentrated, and normalized to 1 mg/mL with PBS supplemented with 1 mM TCEP. And PFO*-EGFP was verified by SDS-PAGE gel.

### PFO*-EGFP staining

1E6 cells were seeded in 12-well plate. On the next day, the cells were treated as required. After treatment, the cells were washed three times with PBS. 1E6 cells were collected and transferred to new tubes for staining. Cell pellet was suspended with 500 µL PBS supplemented with 2% BSA and PFO*-EGFP with the final concentration of 10 µg/mL. The cells were stained at 4°C for 2 h with gentle rotation. After staining, the cells were washed three times with cold PBS and spun down at 500 x g for 5 min at 4°C, followed with fixation with 4% PFA for 10 min at 4°C. The cells were washed and resuspend in 500 µL cold PBS. PFO*-EGFP fluorescence on individual cell was determined on a Fortessa flow cytometer and the MFI was analyzed by FlowJo 10.9.0.

### Confocal microscopy

1E4 of WT and Rabgef1-deficient macrophages were seeded in 12-well chamber (lbidi). On the next day, macrophages were stained with PFO*-EGFP at the final concentration of 10 µg/mL in 200 µL PBS containing 2% BSA for 2 h at 4°C, followed by staining with 0.1 µg /mL Hochest 33342 and 1 µg/mL WGA-Alexa Fluro 647 for 10 min. After staining, macrophages were washed three times with cold PBS and fixed with 4% PFA for 10 min at 4°C, followed with antifade mounting using ProLong Diamond Antifade Mountants without DAPI (Thermo Fisher). The images were captured using Zeiss Celldiscoverer 7. A 405 nM laser was used for Hochest 33342 excitation, a 488 nM laser was used for PFO*-EGFP excitation, a 640 nM laser was used for WGA-Alexa Fluro 647 excitation. PFO*-EGFP intensity was quantified from confocal images using CellProfiler within WGA positive membrane regions constrained by cell outline masks.

### Filipin staining

1E6 of WT and Rabgef1-deficient macrophages were seeded in 12-well plate. On the next day, the cells were washed three times with PBS and lifted. 1E6 cells were fixed with 4% PFA for 10 min at 4°C and stained with filipin at the concentration of 100 µg/mL for 1h at 4°C. Filipin fluorescence was determined on a Fortessa flow cytometer and MFI was analyzed by FlowJo 10.9.0.

### In vitro transcription and mRNA transfection

In vitro transcription of plasmids encoding gasdermins was performed using the HiScribe T7 mRNA Kit with CleanCap Reagent AG (NEB, E2080S) following the manufacturer’s protocol. Briefly, 10 µg plasmids were linearized using Xbal restriction enzyme and purified using QIAquick PCR Purification Kit and eluted with 20 µL nuclease-free water. The concentration was determined by nanodrop. The reaction was assembled at room temperature with the following order:

| Components | 20 μL reaction |
| --- | --- |
| Nuclease-free water | X μL |
| 10X T7 CleanCap Reagent AG Reaction Buffer | 2 μL |
| ATP (60 mM) | 2 μL |
| 5methoxy-UTP (100 mM) | 1 μL |
| 5mCTP (100 mM) | 1 μL |
| GTP (50 mM) | 2 μL |
| Cap Analog (40 mM) | 2 μL |
| Template DNA | 1 μg |
| T7 RNA Polymerase Mix | 2 μL |

After 2 h incubation in thermocycler at 37°C, the IVT reaction was stopped by adding 2 µL DNase I with incubation for 15 min at 37°C. mRNA was purified using the Monarch® Cleanup Kit (500 µg) (NEB, T2050) following the manufacturer’s protocol and the concentration was quantified using nanodrop.

2E5 of macrophages or 1E5 of cancer cells were seeded in 24-well plate. On the next day, the cells were transfected with mRNA encoding gasdermins with EGFP mRNA as complement using Lipofectamine Messenger MAX (Invitrogen, LMRNA015) according to the manufacturer’s protocol. 1 h after transfection, the medium was replaced. Cells were stained with PI at the concentration of 3.3 µg/mL and imaged every 1 h for 8 h using SX5 Incucyte live-cell imager.

### RNA extraction and RT-qPCR

1E6 of WT and Rabgef1-deficient macrophages were seeded in 12-well plate and cultured in complete DMEM or DMEM supplemented with de-lipidated FBS. On the next day, the macrophages were lysed in 1 mL TRIzol and RNA was extracted according to the manufacturer’s protocol. cDNA was synthesized with 2 µg RNA in a 20 µL reaction using High-Capacity cDNA Reverse Transcription kit (Thermo Fisher). RT-qPCR was performed using PowerUp SYBR Green Master Mix (Thermo Fisher). Gene expression was normalized to GAPDH. The primer sequences for mouse GAPDH, SREBP2, LDLR, SQLE and DHCR7 were shown in the oligonucleotides section.

### Mouse Studies

1E6 of Dox inducible B16F10-NT-GSDME-mNeonGreen cells were injected subcutaneously into the right flank of C57BL/6J mice. Tumor growth was monitored every other day by measuring tumor length (a) and width (b) using calipers. Tumor volume was calculated using the formula V= 0.5 x a x b^2^. When tumor volume reached 50-100 mm^3^, mice were randomized into two groups, and standard chow was replaced with Dox chow. Mice received either PBS or AY9944 (25 mg/kg, i.p.) daily for 14 days. Tumor volume and body weight were recorded daily. At the experimental endpoint, tumors were excised, mechanically dissociated and homogenized for flow cytometry analysis. Single cell suspensions were first stained with Fixable Aqua Viability dye (BV510) for 15 min on ice, washed twice with PBS and then stained with anti-CD54 antibody (Alexa Fluor 700) for 30 min on ice, followed by two washes with PBS. Finally, 2E6 stained cells were incubated with PFO*-EGFP for 2 h at 4°C with gentle rotation to assess cholesterol content.

Blood was collected by cheek bleeding using BD Microtainer blood collection tubes. After collection, the tubes were gently inverted to ensure mixing with clot activator, then allowed to stand for 30 min at room temperature to facilitate clot formation. Samples were centrifuged at 3000 x g for 15 min, and the serum was carefully harvested for quantification of total and free cholesterol using Amplex Red Cholesterol Assay kit as described above.

### Statistical analysis

Statistical analysis was performed using GraphPad Prism 10 software. Unpaired two-tailed t-test was used to determine differences between two groups. One-way ANOVA was used to determine differences among multiple groups. Not significant (ns), P > 0.05; *P < 0.05; **P < 0.01; ***P < 0.001; ****P < 0.0001. For western blot analysis, images shown are representative of at least 3 independent experiments. Quantified bar graphs and time course analysis are the combined mean values of three independent experiments, unless otherwise specified in figure legends. Data with error bars are represented as mean ± SEM.

## Acknowledgements

This work was supported by NIH grant GM159938 to C.L.E. C.L.E is also supported by a Ragon Early Independence Fellowship, Karin Grunebaum Cancer Research Foundation Faculty Fellowship, Schwartz Initiative AI/ML Pilot Award, and Ragon Institute Strategic Initiative. Y.L, S.P., C.L.E. and C.S.G. were supported by a Bladder Cancer Advocacy Network Innovation Award 246220 to C.S.G. Y.L is supported in part by the American Heart Association Postdoctoral Fellowship grant 26POST1563608. I.R.E. is a Merck Fellow of the Damon Runyon Cancer Research Foundation, DRG-2508-23. M.H is supported in part by the Japan Society for the Promotion of Science postdoctoral fellowship. G.G.R. is supported in part by the Ramón Areces Foundation Research Fellowship. We thank members of the Evavold lab, Garris lab, Moura Silva lab, Ringel lab, Wong lab, Batista lab, Dang lab, and Lingwood lab for helpful discussions. Biorender was used to generate schematics. We thank Feng Shao for plasmids encoding human gasdermin family members for cloning.

---

**Figure S1. NT-GSDMD oligomerization, but not membrane binding, is defective in Rabgef1 knockout macrophages.**

(A) Cell-Map BioID enrichment analysis of CRISPR-Cas9 screen hits in Dox inducible NT-GSDMD expressing macrophages. (B and C) WT and Rabgef1-deficient macrophages were induced with or without Dox for 8 h. BFP positivity was analyzed by flow cytometry (B) and NT-GSDMD protein expression was determined by western blot (C). (D) Relative TMRM intensity in WT and Rabgef1-deficient macrophages with or without Dox induction for 8 h, normalized to time 0. (E) Western blot analysis of NT-GSDMD from WCL, cytosol and mitochondria fractions in WT and Rabgef1-deficient macrophages after Dox induction for 8 h. Tubulin represents a cytosol marker, and Tomm20 represents a mitochondria marker. Data are mean ± SEM of three repeats. Immunoblots show representative result of three repeats. Statistical significance was determined by one-way ANOVA: *p < 0.05; **p < 0.01; ***p < 0.001; ****p < 0.0001.

**Figure S2. Rabgef1 deficiency has limited effect on internalization pathway in macrophages.**

(A) Schematic of Rabgef1 activities. (B and C) WT and Rabgef1-deficient macrophages were incubated with Zymosan Bioparticles conjugated with Texas Red for 8 h to assess phagocytosis activity. Kinetics of Zymosan Bioparticle positive single cell frequency (B) and intensity (C) were analyzed. (D and E) WT and Rabgef1-deficient macrophages were incubated with Dextran conjugated with Texas Red for 8 h to evaluate pinocytosis activity. Dextran positivity (D) and MFI (E) were analyzed. (F and G) WT and Rabgef1-deficient macrophages were incubated with LDL conjugated with Dil for 8 h to determine the LDLR-mediated endocytosis. LDL positivity (F) and MFI (G) were analyzed. Data are mean ± SEM of three repeats. Statistical significance was determined by one-way ANOVA: *p < 0.05; **p < 0.01; ***p < 0.001; ****p < 0.0001.

**Figure S3. Cholesterol biosynthesis pathway is up-regulated in Rabgef1 deficient macrophages regardless of exogenous lipids.**

(A-D) WT and Rabgef1-deficient macrophages were cultured in complete DMEM or DMEM supplemented with de-lipidated FBS. mRNA levels of LDLR (A), SREBP2 (B), SQLE (C) and DHCR7 (D) were analyzed by RT-qPCR and normalized to GAPDH. (E) Schematic of conjugation of EGFP-SpyCatcher and SpyTag-PFO*. (F) Schematic of PFO*-EGFP binding with plasma membrane cholesterol. (G and H) Coomassie blue staining of EGFP-SpyCatcher and SpyTag-PFO* after purification (G) and conjugation (H). Data are mean ± SEM of three repeats. Statistical significance was determined by unpaired two-tailed t-test: *p < 0.05; **p < 0.01; ***p < 0.001; ****p < 0.0001.

**Figure S4. The physiological cholesterol inhibitor 25-Hydroxycholesterol enforces GSDMD-mediated pyroptosis.**

(A) WT and Rabgef1 sg2 macrophages were treated with 19HC (10 µM) or 25HC (10 µM) for 9 h, followed by PFO*-EGFP staining. The MFI of PFO*-EGFP was analyzed. (B-D) WT and Rabgef1 sg2 macrophages were pre-treated with 19HC (10 µM) or 25HC (10 µM) for 1 h, followed with or without Dox induction for 8 h. Kinetics of PI positive single cell frequency (B), bulk LDH release (C) and cell viability (D) were analyzed. (E) WT and Rabgef1 sg2 macrophages were pre-treated with 19HC (10 µM) or 25HC (10 µM) for 1h, followed with or without Dox induction for indicated time points. NT-GSDMD oligomers and monomers under non-reducing and reducing conditions were analyzed by western blot. Data are mean ± SEM of three repeats. Immunoblots show representative result of three repeats. Statistical significance was determined by one-way ANOVA and unpaired two-tailed t-test: *p < 0.05; **p < 0.01; ***p < 0.001; ****p < 0.0001.

**Figure S5. Rabgef1 deficiency decreases pore formation induced by other gasdermin family members in macrophages.**

(A) Schematic of in vitro transcription (IVT) and mRNA transfection. (B and C) EGFP positive frequency (B) and EGFP intensity (C) in WT and Rabgef1 sg2 macrophages transfected with EGFP mRNA. (D-H) WT and Rabgef1 sg2 macrophages were transfected with mRNA expressing FL and NT of GSDMA3 (D), GSDMB (E), GSDMC (F), GSDMD (G) and GSDME (H). PI positive single cell frequency was analyzed. Data are mean ± SEM of three repeats. Statistical significance was determined by unpaired two-tailed t-test: *p < 0.05; **p < 0.01; ***p < 0.001; ****p < 0.0001.

**Figure S6. Cholesterol depletion sensitizes GSDME mRNA mediated pyroptosis in diverse cancer cell lines.**

(A, E, I and M) Free cholesterol quantification in 1245-PDAC cells (A), LLC1 lung cancer cells (E), MB49 bladder cancer cells (I) and EO771 breast cancer cells (M) treated with or without MCD (7.5 mM) for 1 h. (B, F, J and N) MFI of PFO*-EGFP in 1245-PDAC cells (B), LLC1 lung cancer cells (F), MB49 bladder cancer cells (J) and EO771 breast cancer cells (N) treated with or without MCD (7.5 mM) for 1 h, followed by PFO*-EGFP staining. (C, G, K and O) 1245-PDAC cells (C), LLC1 lung cancer cells (G), MB49 bladder cancer cells (K) and EO771 breast cancer cells (O) were transfected with serial doses of mRNA encoding FL and NT GSDME for 7 h, followed by treated with or without MCD (7.5 mM) for 1 h. PI positive single cell frequency was analyzed. (D, H, L and P) Western blot analysis of FL and NT GSDME expression in 1245-PDAC cells (D), LLC1 cells (H), MB49 cells (L) and EO771 cells (P) transfected with two highest doses of mRNA encoding FL and NT GSDME for 7 h, followed by treatment with or without MCD (7.5 mM) for 1 h. Data are mean ± SEM of three repeats. Statistical significance was determined by unpaired two-tailed t-test: *p < 0.05; **p < 0.01; ***p < 0.001; ****p < 0.0001.

**Figure S7. Reducing cholesterol concentration enhances GSDME pyroptosis in organoids and tumors.**

(A) Western blot analysis of murine NT-GSDME expression in Dox inducible KP pancreatic organoids pre-treated with or without 25HC (10 µM) for 4 h, followed with or without Dox induction for 6 h. (B) Live cell images of Dox inducible KP pancreatic organoids labelled with PI and pre-treated with or without 25HC (10 µM) for 4 h, followed with Dox induction of NT-GSDME expression for 6 h. (C) Kinetics of average PI integrated intensity of Dox inducible KP pancreatic organoids pre-treated with or without 25HC (10 µM) for 4 h, followed with or without Dox induction for 6 h. (D) MFI of PFO*-EGFP from Dox inducible KP pancreatic organoids treated with or without 25HC (10 µM) for 10 h. (E and F) Bulk LDH release (E) and cell viability (F) in Dox inducible KP pancreatic organoids pre-treated with or without 25HC (10 µM) for 4 h, followed with or without Dox induction of NT-GSDME expression for 6 h. (G) Western blot analysis of murine NT-GSDME expression in Dox inducible B16F10-NT-GSDME-mNeonGreen cells pre-treated with 25HC (10 µM) for 8 h, followed with or without Dox induction for 12 h. (H) Live cell images of Dox inducible B16F10-NT-GSDME-mNeonGreen cells labelled with PI and pre-treated with 25HC (10 µM) for 8 h, followed with Dox induction for 12 h. (I) Kinetics of PI positive single cell frequency in Dox inducible B16F10-NT-GSDME-mNeonGreen cells pre-treated with 25HC (10 µM) for 8 h, followed with or without Dox induction for 12h. (J) MFI of PFO*-EGFP in Dox inducible B16F10-NT-GSDME-mNeonGreen cells treated with or without 25HC (10 µM) for 20 h. (K and L) Bulk LDH release (K) and cell viability (L) in Dox inducible B16F10-NT-GSDME-mNeonGreen cells pre-treated with or without 25HC (10 µM) for 8 h, followed with or without Dox induction for 12 h. (M) Melanoma bearing C57BL/6J mice were fed Dox chow and treated daily with AY9944 (25 mg/kg, i.p.) or vehicle for 14 days. Free cholesterol concentration in serum was quantified at the experimental endpoint (n=10). (N) Relative tumor growth in C57BL/6J mice fed Dox chow and treated daily with AY9944 (25 mg/kg, i.p.) or vehicle, normalized to day 0 (n=10). (O) Body weight of C57BL/6J mice fed Dox chow and treated daily with AY9944 (25 mg/kg, i.p.) or vehicle over time (n=10). For C-F, data are mean ± SEM of four repeats. For I-L, data are mean ± SEM of three repeats. Immunoblots show representative result of at least three repeats. For M-O, data are mean ± SEM and are representative of two repeats. Statistical significance was determined by unpaired two-tailed t-test: *p < 0.05; **p < 0.01; ***p < 0.001; ****p < 0.0001.

**Figure S8. Chemotherapy induced GSDME pyroptosis is enhanced by cholesterol depletion.**

(A and B) mNeonGreen and FL-GSDME-mNeonGreen stably expressing B16F10 cells were pre-treated with or without AY9944 (10 µM) (A) or 25HC (10 µM) (B) for 8 h, followed by treatment with or without Raptinal (5 µM) for 2 h. Expression and cleavage of FL-GSDME were analyzed by western blot. (C and E) mNeonGreen and FL-GSDME-mNeonGreen stably expressing B16F10 cells were pre-treated with or without AY9944 (10 µM) for 8 h, followed by treatment with or without Raptinal (5 µM) for 2 h. PI positive single cell frequency (C), bulk LDH release (D) and cell viability (E) were analyzed. (F and H) mNeonGreen and FL-GSDME-mNeonGreen stably expressing B16F10 cells were pre-treated with or without 25HC (10 µM) for 8 h, followed by treatment with or without Raptinal (5 µM) for 2 h. PI positive single cell frequency (F), bulk LDH release (G) and cell viability (H) were analyzed. (I and J) mNeonGreen and FL-GSDME-mNeonGreen stably expressing B16F10 cells were pre-treated with or without AY9944 (10 µM) (I) or 25HC (10 µM) (J) for 8 h, followed by treatment with or without Cisplatin (160 µM) for 16 h. Expression and cleavage of FL-GSDME were analyzed by western blot. (K and M) mNeonGreen and FL-GSDME-mNeonGreen stably expressing B16F10 cells were pre-treated with or without AY9944 (10 µM) for 8 h, followed by treatment with or without Cisplatin (160 µM) for 16 h. PI positive single cell frequency (K), bulk LDH release (L) and cell viability (M) were analyzed. (N and P) mNeonGreen and FL-GSDME-mNeonGreen stably expressing B16F10 cells were pre-treated with or without 25HC (10 µM) for 8 h, followed by treatment with or without Cisplatin (160 µM) for 16 h. PI positive single cell frequency (N), bulk LDH release (O) and cell viability (P) were analyzed. Data are mean ± SEM of four repeats. Statistical significance was determined by unpaired two-tailed t-test: *p < 0.05; **p < 0.01; ***p < 0.001; ****p < 0.0001.

## Notes

### Competing Interest Statement

The authors have declared no competing interest.

