## Supplemental_Figures for "Cholesterol constrains pore formation by mammalian gasdermins"

Figure S1. NT-GSDMD oligomerization, but not membrane binding, is defective in Rabgef1 knockout macrophages.

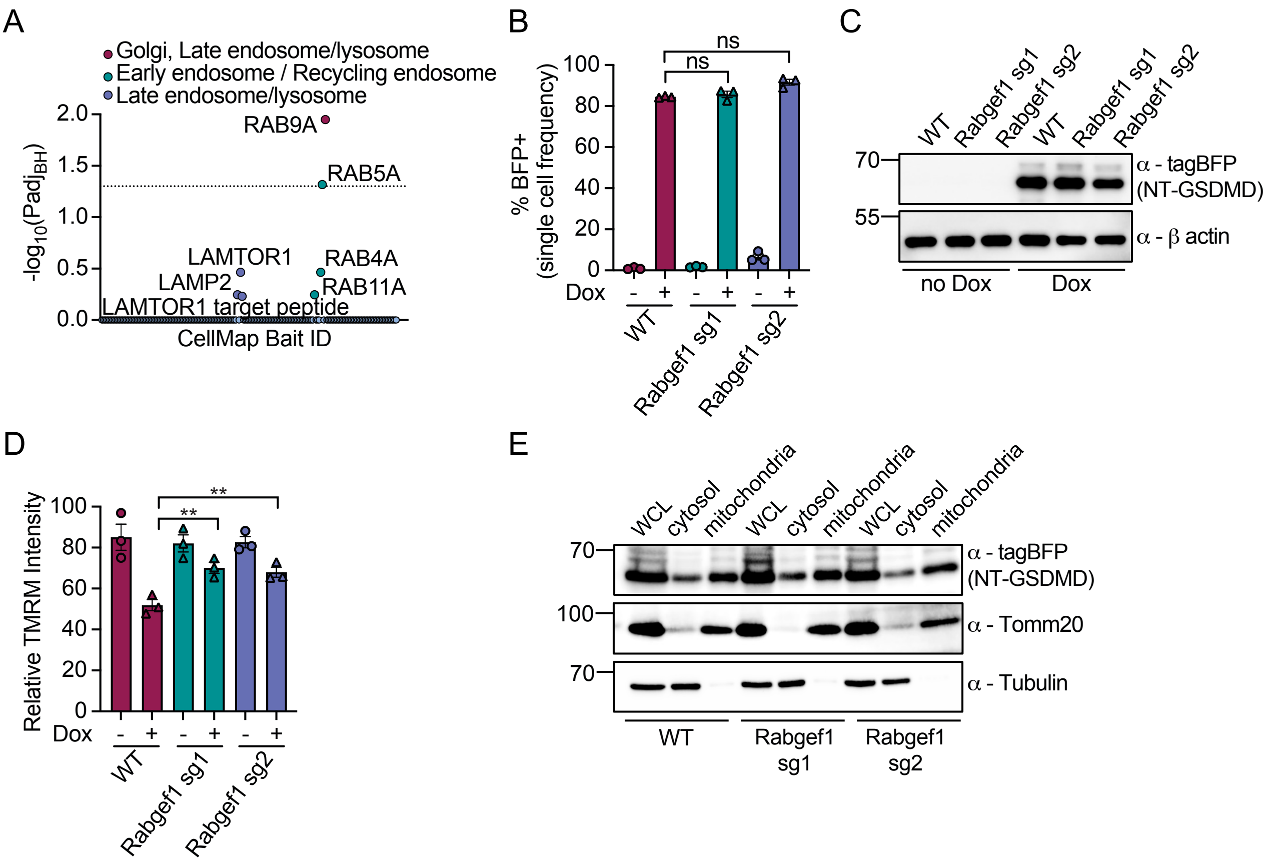

Figure S2. Rabgef1 deficiency has limited effect on internalization pathways in macrophages.

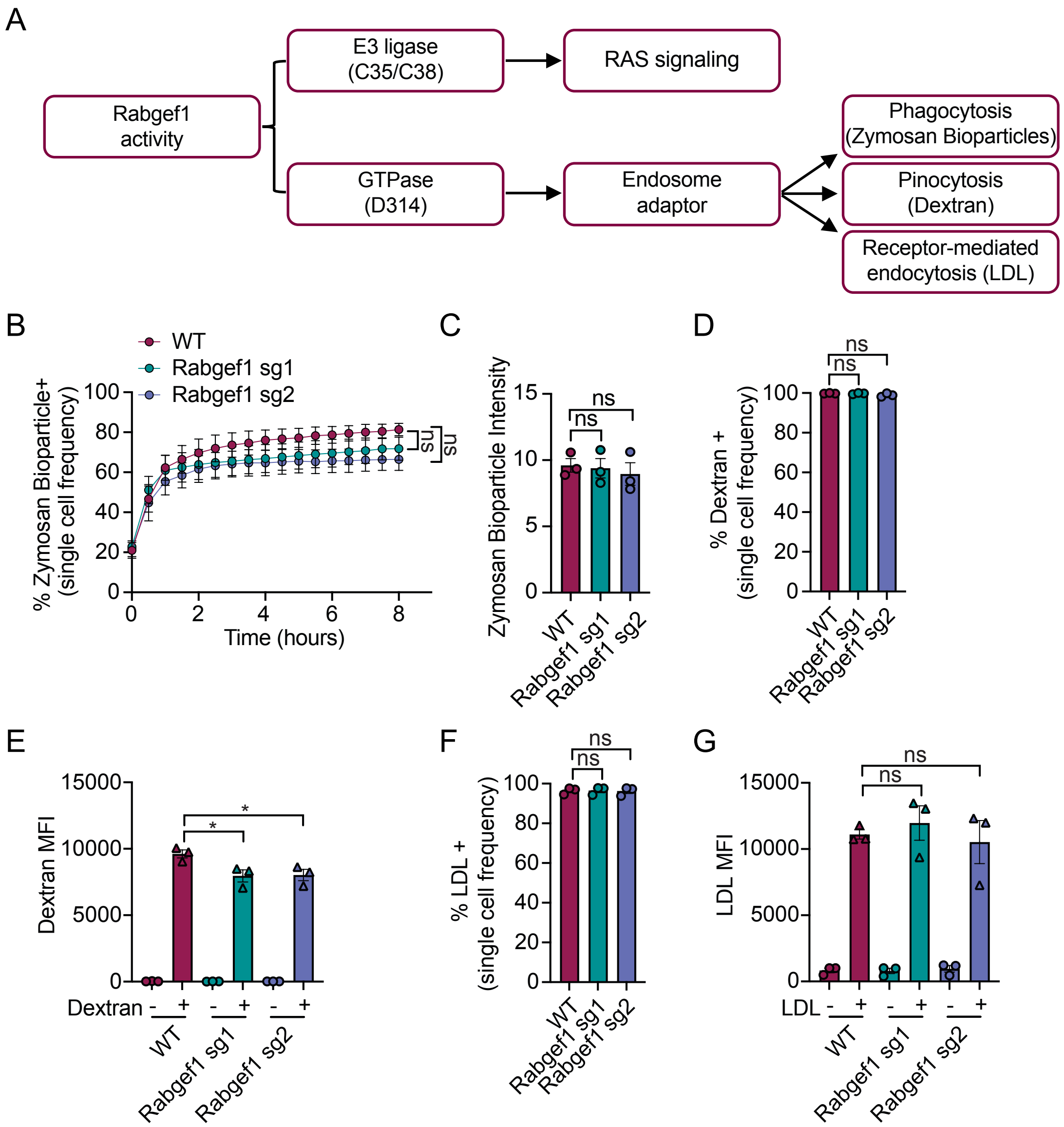

Figure S3. Cholesterol biosynthesis pathway is up-regulated in Rabgef1 deficient macrophages regardless of exogenous lipids.

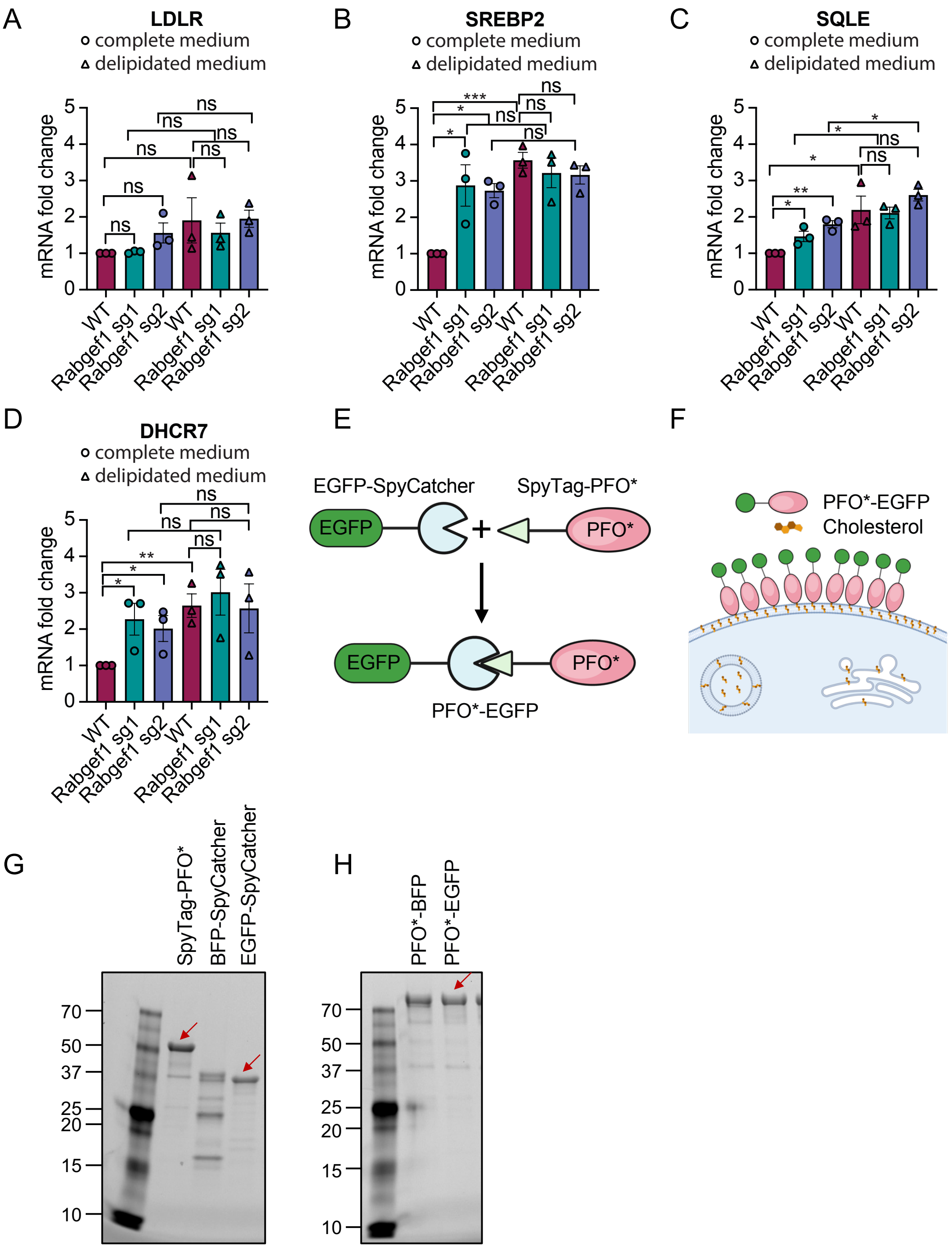

Figure S4. The physiological cholesterol inhibitor 25-Hydroxycholesterol enforces GSDMD-mediated pyroptosis.

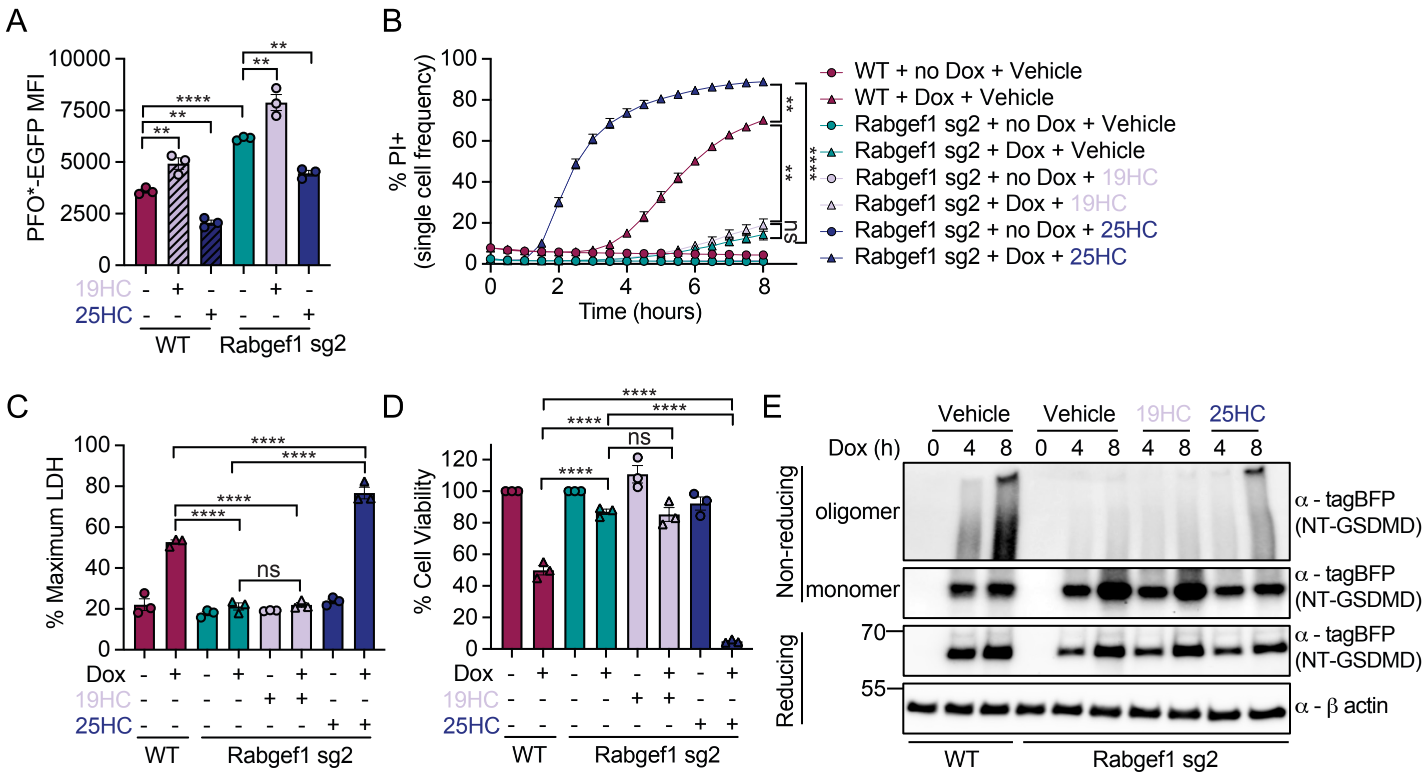

Figure S5. Rabgef1 deficiency decreases pore formation induced by other gasdermin family members in macrophages.

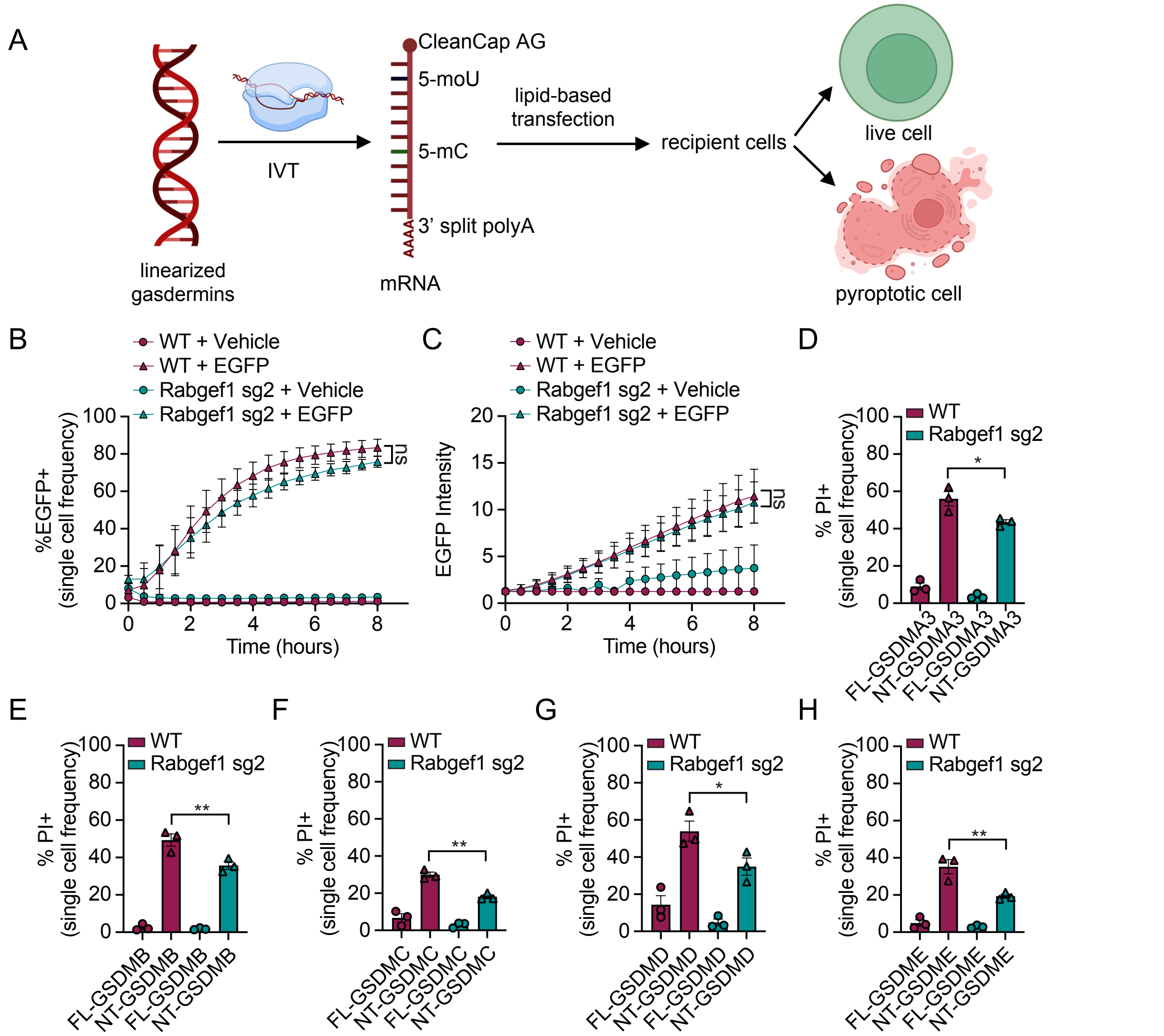

Figure S6. Cholesterol depletion sensitizes GSDME mRNA mediated pyroptosis in diverse cancer cell lines.

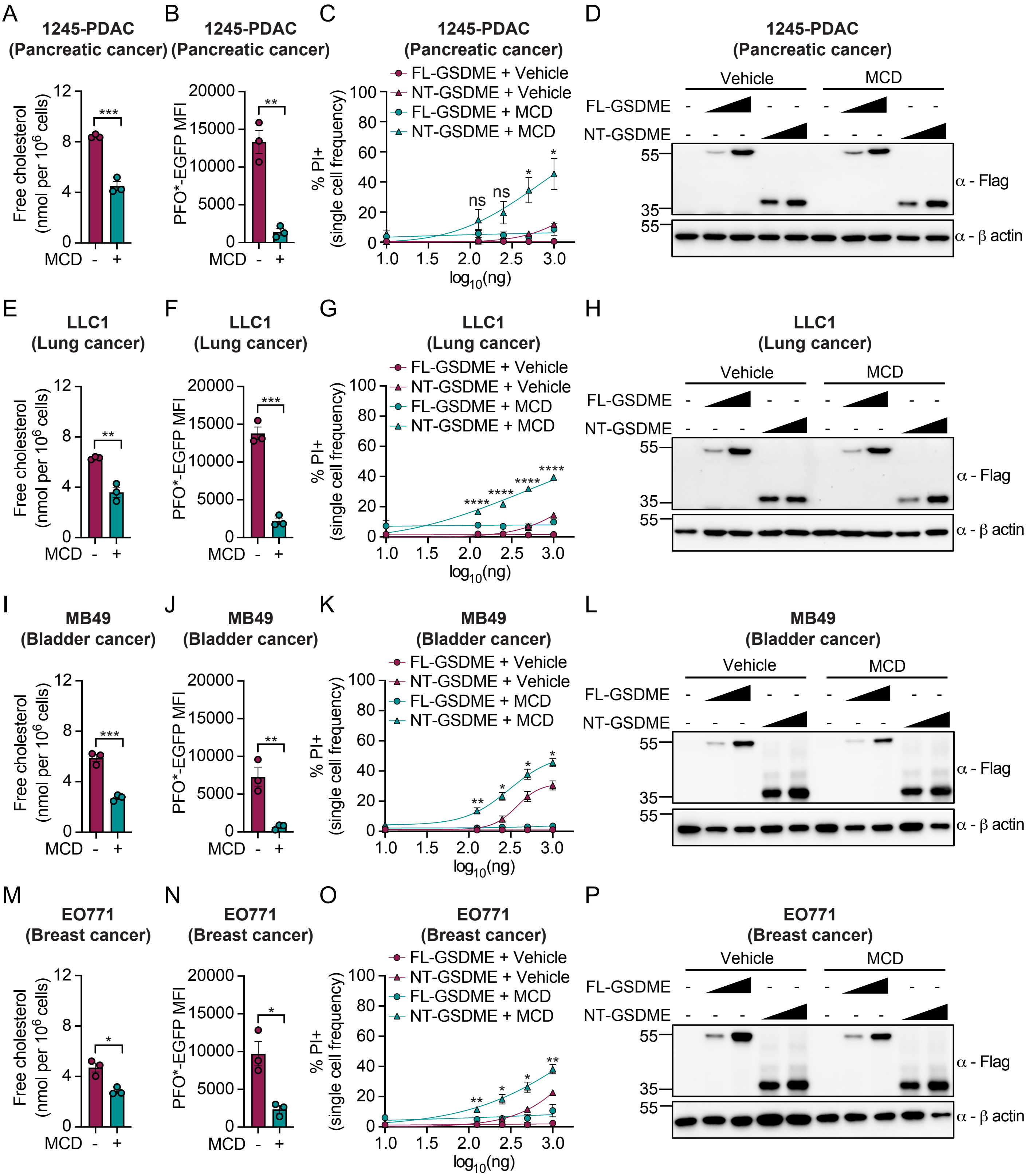

Figure S7. Reducing cholesterol concentration enhances GSDME pyroptosis in organoids and tumors.

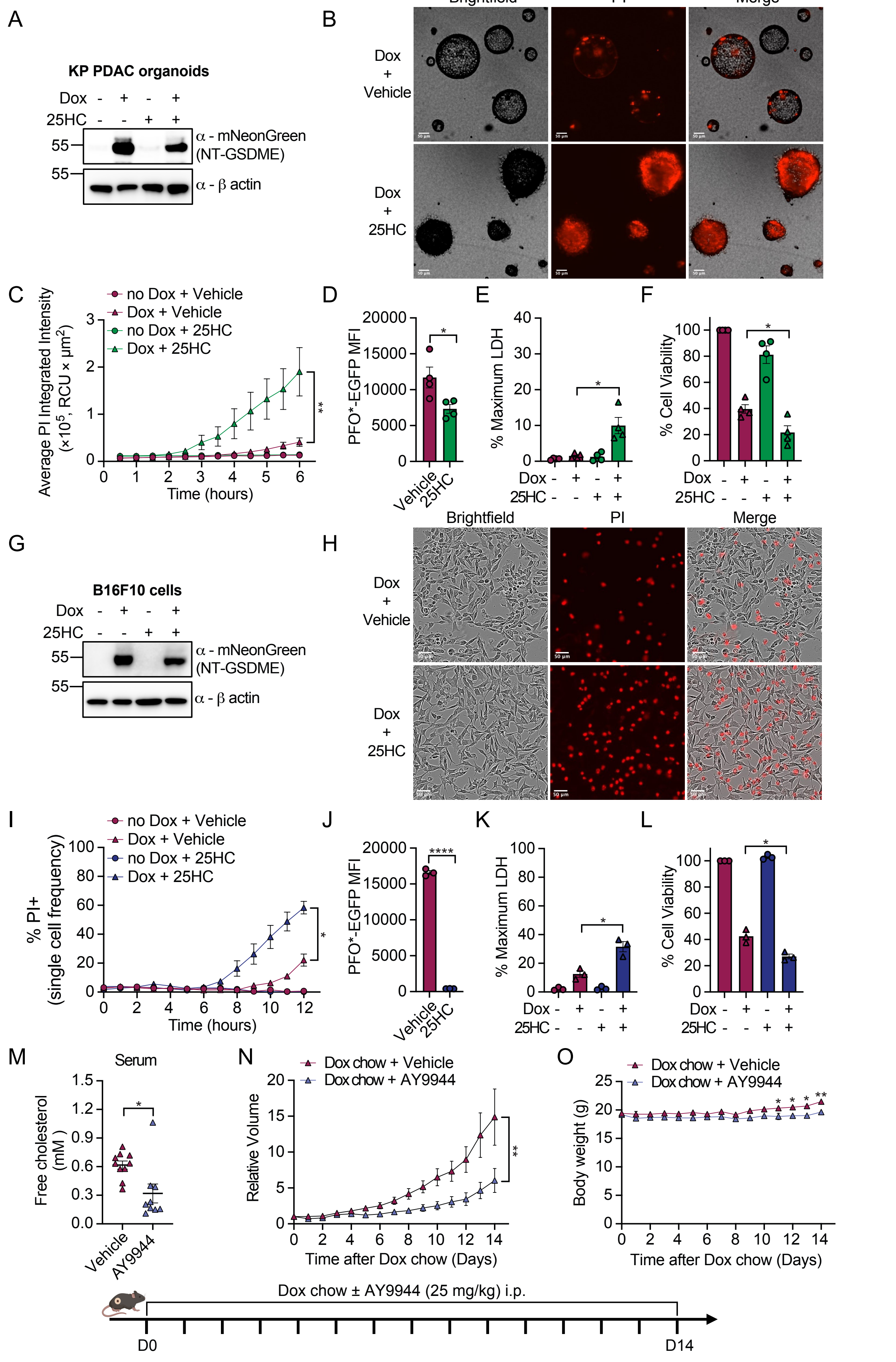

Figure S8. Chemotherapy induced GSDME pyroptosis is enhanced by cholesterol depletion.

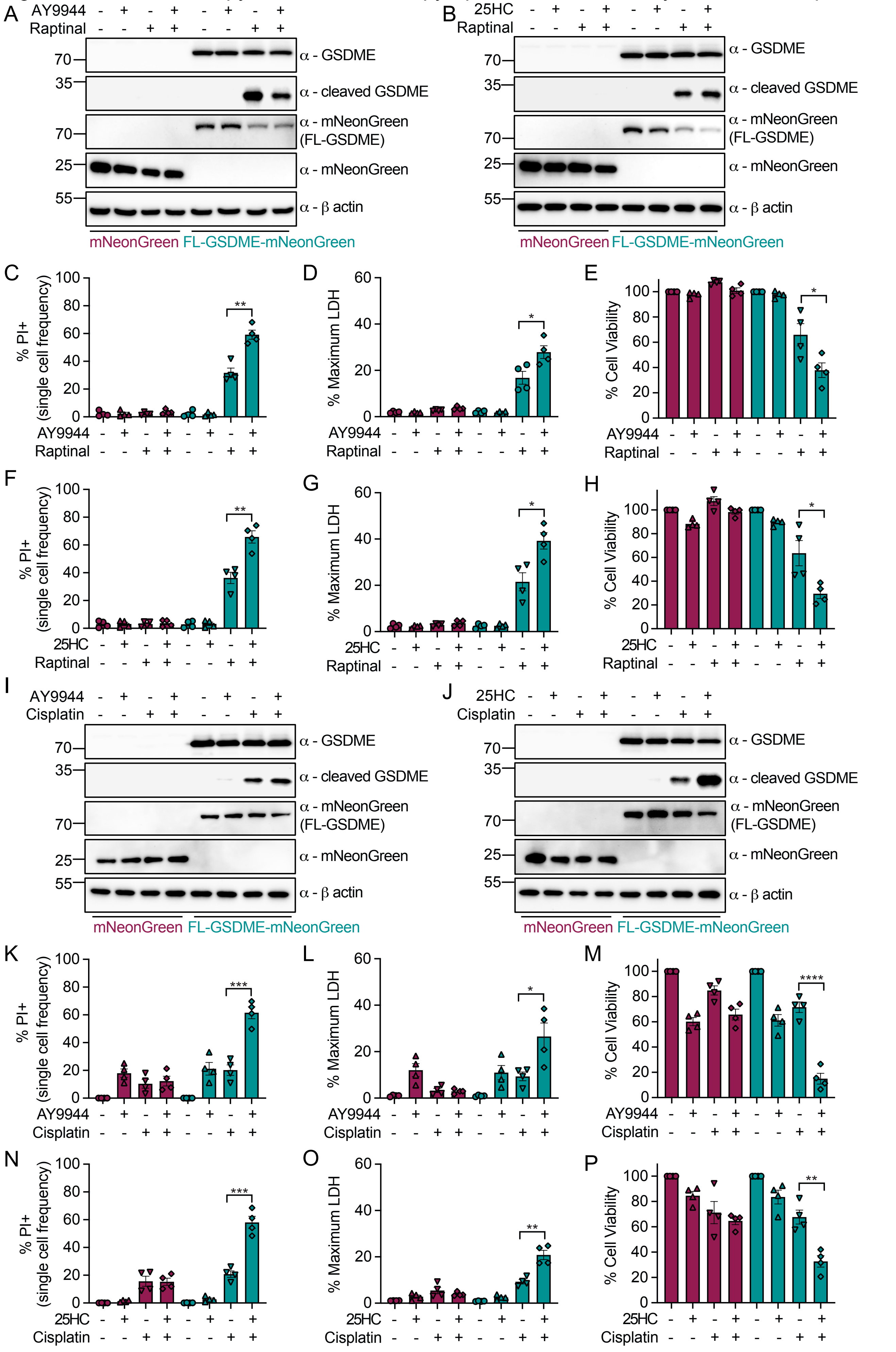
